# Microglia surveillance is directed toward neuron activation during sustained intracortical microstimulation

**DOI:** 10.1101/2025.06.09.658722

**Authors:** C Preszler, K Stieger, K Chen, G Zhang, TDY Kozai

**Affiliations:** Department of Bioengineering, University of Pittsburgh, USA; Center for the Neural Basis of Cognition, USA; Center for Neuroscience, University of Pittsburgh, USA; McGowan Institute for Regenerative Medicine, University of Pittsburgh, Pittsburgh, PA, USA; Neuroscience Institute, Carnegie Mellon University, Pittsburgh, PA, USA

**Author notes:** = denotes equal contributions (C Preszler, K Stieger, K Chen).

**Keywords:** Gliomodulation, immunoelectrophysiology, antidromic activation, gliosynaptic transmission, microglia-neuron crosstalk, neural hysteresis, tripartite synaptic modulation, neuroimmune feedback

## Abstract

Intracortical microstimulation (ICMS) is a widely used tool for neuroprostheses, but its long-term efficacy is often limited by biofouling and neuroinflammatory responses at the electrode-tissue interface. Microglia orchestrate neuroinflammation and regulate synaptic plasticity, and low-frequency stimulation has been shown to promote anti-inflammatory microglial phenotypes. We therefore examined, over the first three days post-implantation, how 10-Hz ICMS influences microglial-neuronal interactions in vivo using two-photon imaging in Cx3cr1-GFP/jRGECO1a mice.

A one-hour session of 10-Hz ICMS did not induce overt morphological activation of microglia but increased their process motility, directing extensions toward both the electrode and neurons exhibiting elevated calcium activity. By post-implantation Day 2, microglial extensions were significantly biased toward neurons whose ΔF/F surpassed a 3 standard deviation threshold after stimulation onset (74.26° ± 11.83°) but shifted away from those same neurons after 40 min of continuous stimulation (116.99° ± 9.19°) (p = 0.001732), suggesting a dynamic, homeostatic response to sustained neuronal calcium elevations. Although multi-day electrode insertion accelerated microglial motility and polarization toward the device, 10-Hz ICMS alone did not alter microglial branching or soma shape. Microglial contact frequency scaled with neuronal adaptation profiles: depressed neurons received the most contacts immediately post-implant (1.15 ± 0.3 contacts; p = .0460). These findings reveal stimulus-associated, neuron-dependent surveillance behaviors of microglia during early post-implantation ICMS and implicate them as active participants in short-term modulation of cortical circuits.

## Introduction

Electrical stimulation is a cornerstone of neurotechnology, enabling therapeutic neuromodulation and bidirectional communication with neural circuits via implanted devices [1–6]. However, long-term efficacy is often limited by biofouling and chronic neuroinflammatory responses triggered by electrode-tissue interfaces [1, 3, 7–13]. While neuronal responses to intracortical microstimulation (ICMS) are well-characterized [5, 14–16], the contribution of microglia, the brain’s primary immune cells, to stimulation outcomes remains poorly understood [17–21].

Microglia are not merely passive responders to injury, they dynamically regulate synaptic plasticity, neuroinflammation, and electrode biocompatibility through cytokine signaling (e.g., TNF-α, IL-1β) and the formation of physical barriers in brain tissue [22–31]. Electrical stimulation through conductive materials may influence these microglia-neuron interactions [2, 5, 6, 15, 16, 32–35] thereby potentially modulating glia-neuron signaling and impacting the fidelity and longevity of neuromodulation[36].

Strategies to mitigate microglial activation have focused on surgical technique [37–40], pharmacological modulation[41–45], and electrode surface engineering [23, 25, 46–48]. Several studies support the notion that the increase in trophic factor release (e.g. BDNF) during 2-20 Hz ICMS leads to the decrease of pro-inflammatory cytokines from microglia, increase in angiogenesis, and improvement in outcome measures[2, 49, 50]; however, the potential of electrical stimulation itself to engage or regulate microglia-neuron dynamics is underexplored [36]. Specifically, it is unclear whether microglia respond to ongoing electrical stimulation in vivo and if it could shift microglia toward or away from an inflammatory phenotype. Given that microglial activation states may govern the balance between functional integration and fibrotic encapsulation[20, 21, 51–53], a deeper understanding of how ICMS modulates microglial interactions with neurons is essential for optimizing stimulation paradigms and improving performance of neuroprostheses.

Microglia behavior and physiology may be directly influenced by electrical fields, independent of neuronal activity, due to sensitivity to membrane polarization and field-induced ion flux. Although they do not generate action potentials, microglia express voltage-gated ion channels, purinergic receptors, and electrochemically sensitive pathways [44, 54–56]. Electric fields can perturb extracellular ion concentrations (e.g., Ca²⁺, K⁺) or redistribute charge across the microglial membrane, activating downstream pathways that mediate pro-inflammatory cytokine release (e.g., via NF-κB) or anti-inflammatory signaling (e.g., via PPAR-γ) [54, 57–61]. Notably, *in vitro* studies demonstrate that microglia exhibit directed migration, increased process motility, and morphological remodeling in response to direct electric fields [23, 62–67]. These findings suggest that specific stimulation parameters could be engineered to modulate their behavior. For neuroprostheses, this raises a provocative possibility: electrical stimulation might serve as an intervention to promote biocompatibility by actively suppressing detrimental biofouling (e.g., cytokine-mediated fibrosis) while promoting tissue-electrode integration.

Following device implantation, microglia transition to a reactive state within minutes by extending processes toward the implant site [43, 44, 68, 69] followed by the release of pro-inflammatory cytokines such as TNF-α and IL-1β within hours [15, 23, 70–72]. These cytokines exacerbate oxidative stress and neuronal damage while also driving glial scar formation [28, 31], which increases interfacial impedance and degrades signal fidelity [9, 11–13, 41, 73–84]. Microglia also mediate synaptic stripping and aberrant plasticity near electrodes [46, 63, 85–88], potentially impairing therapeutic stimulation.

However, microglia also clear cellular debris and secrete neurotrophic factors [18, 86] indicating a context-dependent role that could be harnessed to improve device integration. For example, fractalkine signaling (CX3CL1-CX3CR1) downregulates excessive microglial reactivity [72, 89–91], while adenosine (a byproduct of microglial ATP metabolism) suppresses neuronal hyperexcitability [61, 92–94]. In neuroprosthetic applications, this suggests a strategy wherein appropriately tuned electrical stimulation could modulate microglial states to attenuate cytokine-driven inflammation to promote microglia-mediated tissue integration at the electrode interface.

Here, we investigate how 1 hour of 10-Hz ICMS influences microglial-neuronal interactions *in vivo*, using two-photon imaging to quantify neuronal calcium activity and microglia process dynamics in mice expressing GFP in microglia and jRGECO1a in Thy1+ neurons (CX3CR1-GFP x Thy1-jRGECO1a). We focus on low-frequency stimulation due to previous studies demonstrating reduced inflammatory states in microglia [49, 50, 95]. Additionally, since microglia have been demonstrated to suppress neural activity [96], this animal model provided the opportunity to investigate the relationship between microglia behavior (e.g. putative contacts with neurons) and reduction in neural activity during continuous stimulation. Our results show that prolonged ICMS induces microglial process extension toward both the electrode site and surrounding activated neurons, with microglia more frequently contacting neurons that exhibit stronger adaptation in calcium activity. These interactions suggest low-frequency ICMS facilitates a functional coupling in which microglia may preferentially surveil or stabilize active neurons. These results position microglia as dynamic, neuron-responsive elements within a stimulated circuit and reveal stimulation-dependent patterns of microglial-neuronal interaction. By identifying activity-dependent microglial responses to ICMS, this study provides a framework for probing how glial dynamics contribute to the efficacy and stability of electrical neuromodulation.

## Methods

### 2.1 Experimental animal models

All animal care and procedures were performed with the approval of the University of Pittsburgh Institutional Animal Care and Use Committee and in accordance with regulations specified by the Division of Laboratory Animal Resources. Mice were kept under standard conditions on a 12 h light/ dark cycle with access to water and food *ad libitum*. B6.129P2(Cg)-Cx3cr1^tm1Litt^/J (Cx3cr1^GFP^, strain# 5582, Jackson Laboratories; Bar Harbor, ME) were bred with Tg(Thy1-jRGECO1a)GP8.62Dkim/J (jRGECO1a, strain# 30528, Jackson Laboratories; Bar Harbor, ME) to produce Cx3cr1^GFP^/jRGECO1a mice, permitting quantification of microglia morphology (GFP) and neuronal calcium activity (jRGECO1a) (Figure 1a).

**Figure 1.**
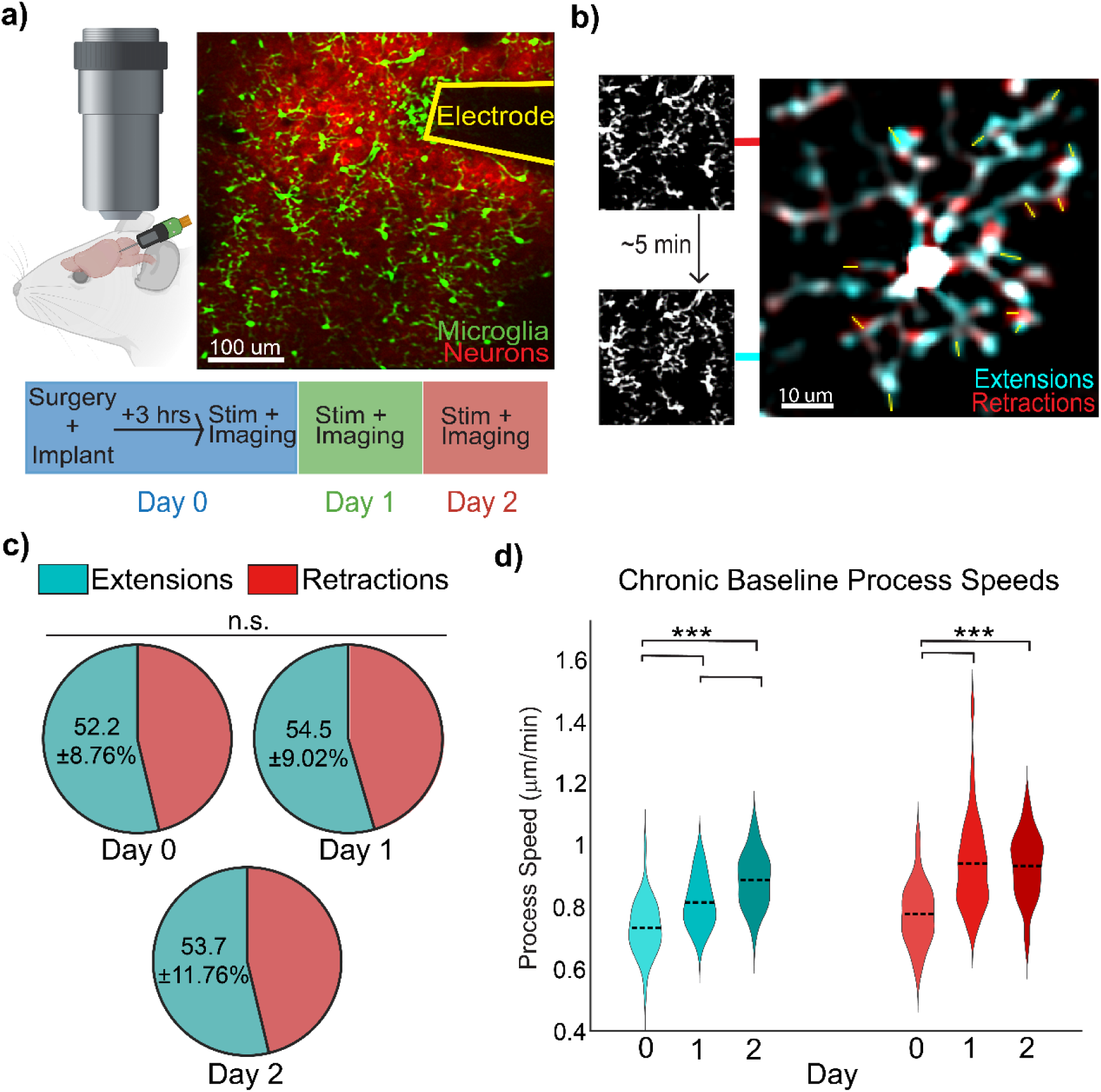
Chronic microelectrode insertion increases the speed of microglial process movements. a) Experimental setup for two-photon microscopy of Cx3CR1-GFP/Thy1-jRGECO1a mice implanted with a single-shank microelectrode in L2/3 of the visual cortex. b) Representative analysis of microglial process motility using FIJI. Time-lapsed images were overlayed in 5 min intervals to highlight extensions (blue) and retractions (red). c) The net balance of microglial process movements (extension vs. retraction) remains unchanged following chronic microelectrode insertion (N=4 mice, n=39,38,37 microglia, One-Way ANOVA, p=0.591). d) Chronic electrode implantation significantly increases the rate of both process extension and retraction in microglia (N=4 mice, n=39,38,37 microglia, One-Way ANOVA, p = 1.06e-4,3.45e-12,9.1e-4 for extensions and p = 9.59e-8,4.21e-7,0.957 for retractions).

**Figure 2.**
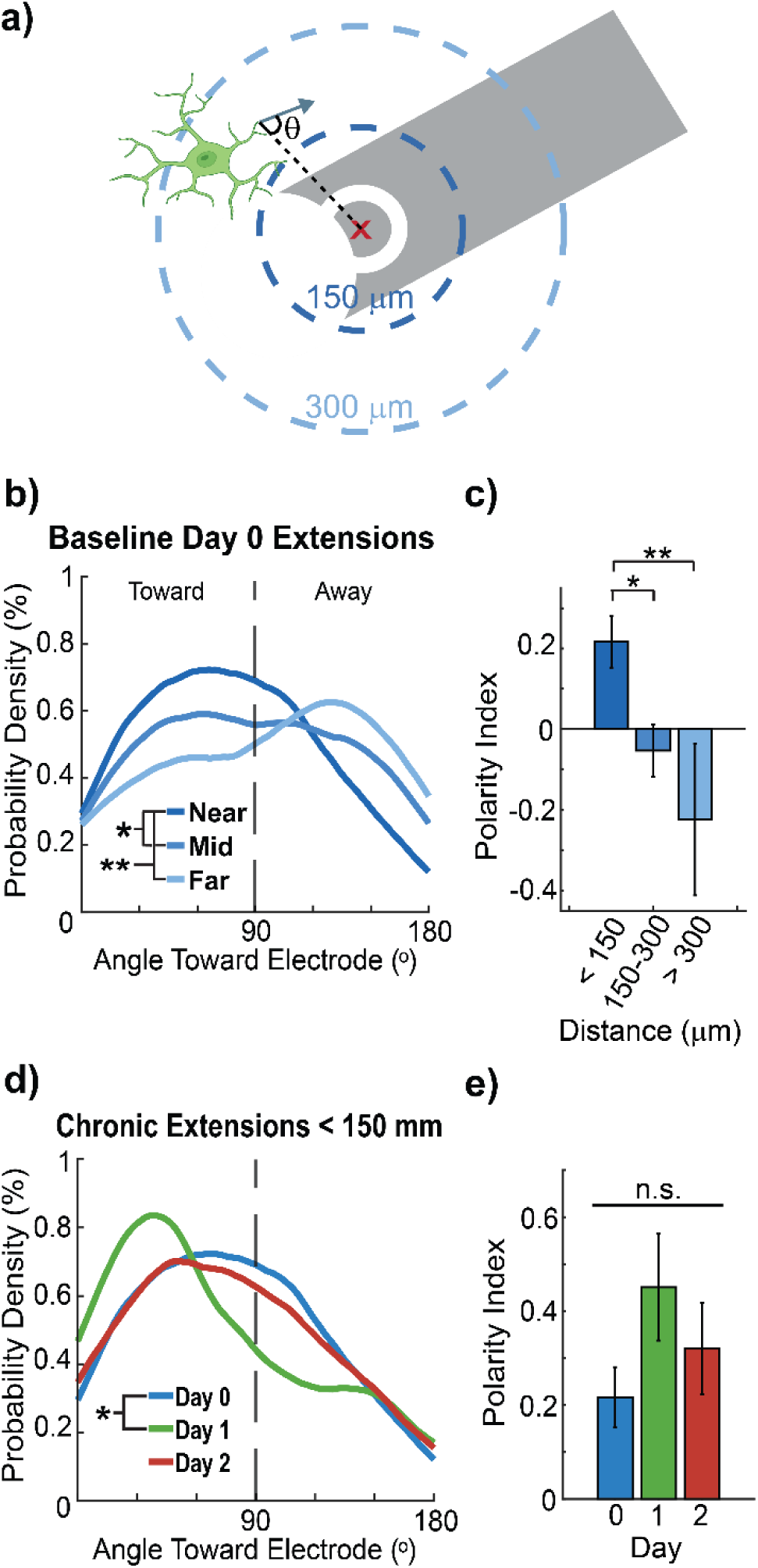
Microglia within 150 µm of chronic microelectrodes extend processes more frequently toward the electrode. a) Measuring the directionality of microglia process extensions relative to the stimulation site. b) The distribution of microglia extensions toward the electrode is inversely proportional to the distance from the electrode (Kolmogorov-Smirnov, p = 0.043, .002, .292). c) Microglia extensions are more polarized toward the electrode within 150 um (One-way ANOVA, p = .031, .021, .776, n = 18, 27, 6 microglia with processes identified in the distance grouping, microglia were excluded if there were fewer than 4 movements). d) Microglia within 150 µm of the electrode peak in their distribution of process extensions toward the electrode on Day 1 (Kolmogorov-Smirnov, p = .034, 1, .118). e) Extensions of microglia processes within 150 µm of the microelectrode do not become more polarized chronically (One-way ANOVA, p = .172, n=18,10,11 microglia, microglia with fewer than 4 movements were excluded from p-index calculations).

### 2.2 Probe implantation surgery

Cx3cr1^GFP^/jRGECO1a mice (n= 2 male, 2 female, < 6 mo., 25-35 g) were implanted with a single-shank Michigan style microelectrode array for awake, head-fixed imaging, as described previously [9, 11, 74, 97–99]. The mice were sedated with a cocktail of 7 mg/kg and 75 mg/kg ketamine prior to removal of the sterilized scalp and drilling of bilateral craniotomies over the visual cortices. Electrodes were targeted to the right hemisphere unless there was bleeding or injury. Prior to implantation, iridium electrode sites were activated to increase charge storage capacity [100] and 1kHz impedance was verified to be <500 kOhm. Bone screws were implanted over the motor cortices for stability and for ground and reference. Single-shank 3 mm long electrode arrays with 4 Iridium channels of 702 µm^2^ electrode sites spaced 50 µm apart (NeuroNexus Technologies, Ann Arbor, MI) were implanted at a 30° angle from the horizontal plane using a Microdrive (MO-81, Narishige, Japan) so that stimulation occurred in layer 2/3 of the visual cortex (final depth of 200-300 µm below the surface). Once the electrode was inserted, the craniotomies were filled with sealant (Kwik-Sil) before being sealed with glass coverslips and dental cement. Ketofen (5 mg/kg) was provided post-operatively up to two days post-surgery or as needed. Animals were allowed to wake up fully prior to any stimulation and imaging (3+ hours).

### 2.3 Stimulation Paradigm

Stimulation was provided using a TDT IZ2 stimulator controlled by an RZ5D system (Tucker-Davis Technologies, Alachua, FL). The stimulation paradigm is illustrated in Figure 3a and is biphasic, symmetric, and cathodic-leading with 200 µs pulse width and 10 µA resulting in 2 nC/phase charge injection, well below the 4 nC/phase safety limit to ensure maximal biocompatibility [7, 101] even for a stimulation trial lasting an hour. Stimulation trials included a 20-min baseline followed by a 60-min stimulation period and a 20-min post-stimulation period (Figure 3a).

**Figure 3.**
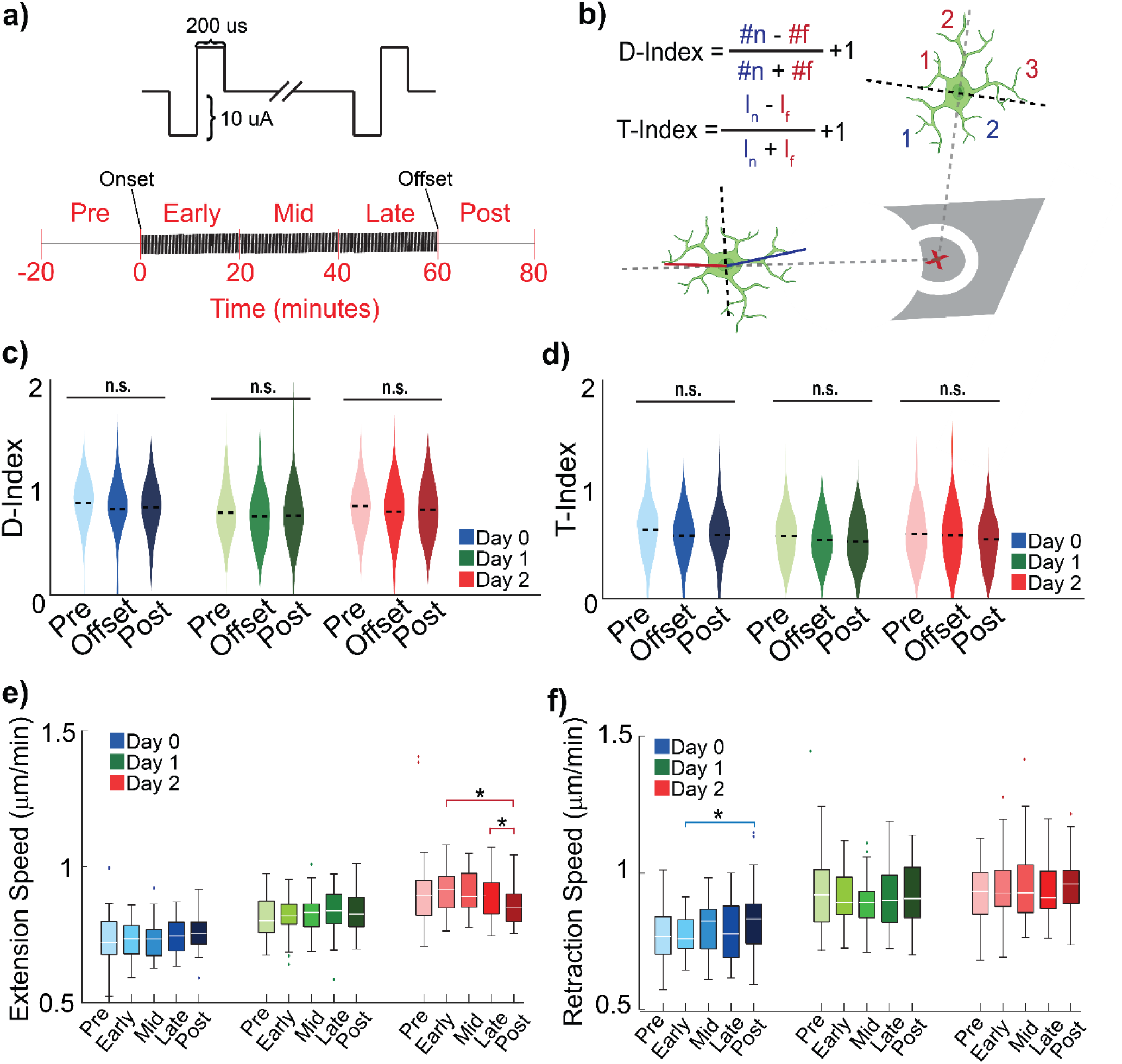
10-Hz ICMS does not affect nearby microglia morphology but modulates process movement speeds. a) ICMS paradigm for 10-Hz symmetric, cathodic-leading, biphasic pulses with a pulse width of 200 µs for 1 h. b) D- and T-indices represent microglia morphology and activation, quantified using the direction and length of the processes, respectively. c) One hour of 10-Hz ICMS did not significantly modulate the directionality of microglia processes and thus does not modulate morphological indicators of microglia activation (Repeated Measures ANOVA, p = 0.296, 0.256, 0.417). d) 10-Hz ICMS did not significantly modulate the length of microglia processes relative to the electrode and thus does not modulate another morphological indicator of microglia activation (Repeated Measures ANOVA, p = 0.132, 0.570, 0.329). e) The offset of 10-Hz ICMS on Day 2 reduced microglia extension speeds (Repeated Measures ANOVA, p = 0.038, 0.036). f) The offset of 10-Hz ICMS on Day 0 increased microglia retraction speeds (Repeated Measures ANOVA, p = 0.042).

**Figure 4.**
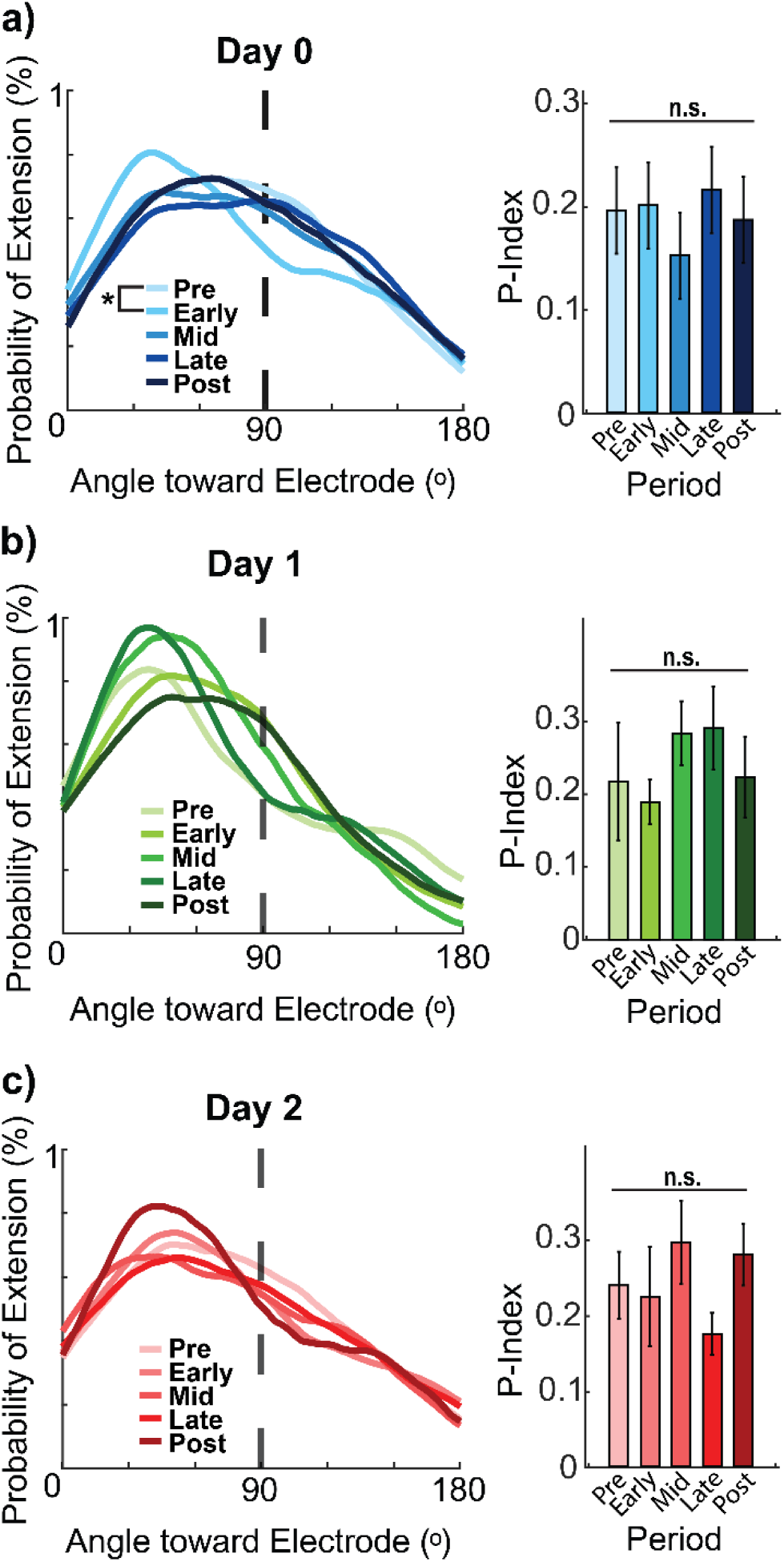
10-Hz ICMS drives microglia to extend processes more frequently toward the electrode only on Day 0. a) On Day 0, the onset of ICMS drives microglia processes within 150 μm of the electrode to extend more frequently toward the electrode but does not significantly increase polarization (N = 39 microglia, n = 194, 174, 187, 187, 165 process extensions, Kolmogorov-Smirnov, p=.026 and Repeated Measures ANOVA, p = .865). b) By Day 1, microglia process extensions within 150 µm are not significantly driven toward the electrode, nor polarized by stimulation (N = 38 microglia, n = 106, 108, 96, 96, 73 process extensions, Kolmogorov-Smirnov, p = .324 and Repeated Measures ANOVA, p = .228). c) Similarly, on Day 2, ICMS does not significantly drive microglia extensions toward the electrode, nor affect polarization (N = 37 microglia, n = 132, 71, 88, 99, 93 process extensions, Kolmogorov-Smirnov, p > .21 and Repeated Measures ANOVA, p = .293).

**Figure 5.**
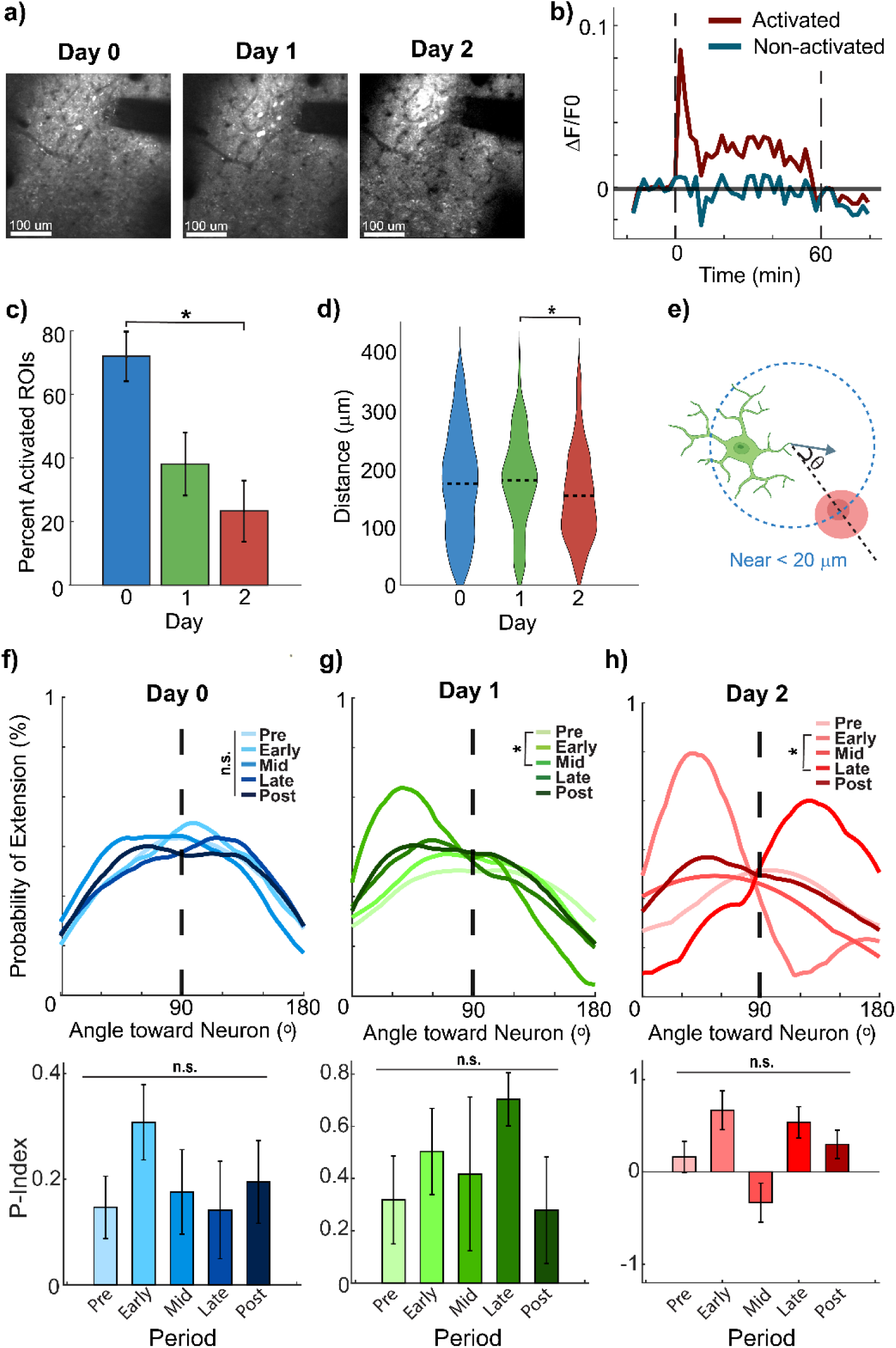
Neuronal activation decreases with chronic 10-Hz ICMS while microglia begin to extend processes more frequently toward activated neurons by Day 1. a) Representative chronic images of activated neurons (Thy1-jRGECO1a). b) Representative calcium traces of neurons, defined as activated and non-activated. c) The percentage of activated neurons decreases significantly between Days 0 and 2 (N = 4 mice, One-Way ANOVA, p = .013). d) Neuron activation shifts closer to the electrode site by Day 2 (N = 4 mice, n = 335, 140, 83 neurons, One-Way ANOVA, p = .040). e) Microglia process extensions within 20 µm of the nearest neuron were considered nearby. f) Microglia process extensions on Day 0 are not directed toward nearby activated neurons regardless of stimulation (N = 39 microglia, n = 214, 184, 159, 193, 177 process extensions Kolmogorov-Smirnov, p > .27, Repeated Measure ANOVA, p = .234). g) On Day 1, microglia process extensions are more directed toward activated neurons after 20 min of 10-Hz ICMS but are not significantly polarized (N = 38 microglia, n = 52, 44, 48, 40, 42 process extensions, Kolmogorov-Smirnov, p > .27) but not significantly polarized (Repeated Measure ANOVA, p = .149). h) On Day 2, microglia process extensions are more directed toward nearby activated neurons at the stimulation onset but shift away from activated neurons after 40 min of stimulation (N = 37 microglia, n = 6, 13, 3, 12, 9 process extensions, Kolmogorov-Smirnov, p = .002). Microglia are not significantly polarized toward activated neurons during stimulation (Repeated Measure ANOVA, p = .313).

### 2.4 Two-photon imaging and stimulation

Imaging of neurons and microglia were achieved with a two-photon microscope (Bruker, Madison, WI) with an OPO laser (Insight DS+, Spectra Physics, Menlo Park, CA) equipped with a 16X 0.8 NA water immersion objective (Nikon Instruments, Melville, NY) with a 3mm working distance resulting FOV of 407 x 407 µm^2^ (1024 x 1024 pixels). Image acquisition consisted of ZT-series with a 2-µm step size over a 20 µm stack size with a 4.6µs dwell time resulting in a stack period of 60.4517s. The wavelength of the latter alternated between stacks, starting with 1060nm to image neuron activity, and 920 nm to image microglia morphology. While this imaging approach reduced temporal resolution, it minimized microglial process displacement across planes, maintained high signal-to-noise ratios for both cell types, and limited thermal loading near the electrode. Imaging time points were 3 h, 24 h, and 48 h post-implantation to assess acute and early chronic responses. Timing metadata for each stack was exported for later synchronization with other datasets.

### 2.5 Data Analysis

#### 2.5.1 Microglia analysis

The microglia ZT-series was processed using a Richardson-Lucy Total Variation Deconvolution in ImageJ [102] followed by 3D Gaussian blurring and background subtraction. Following rigid motion correction, the series was then condensed to a 2D plane by taking the average of each Z-stack to capture as many microglia processes as possible regardless of their direction. The process motility was manually measured in FIJI [103] via overlayed scans taken 4.32-min (2 frames) apart, like previous studies [104], where cyan represents extensions of processes and red represents retractions (Figure 1b). To avoid quantifying process movements out of the imaging volume, microglia were included only if they exhibited typical indicators of surveillance [17, 105], including repeated extensions and retractions. Measurements of process movements were verified by viewing the original T-series. Only process movements were captured during these measurements and were manually classified as extensions or retractions.

The properties of these process movements were calculated using MATLAB, and the angle or directionality of the process movements were calculated relative to either the electrode site or the nearest neuronal soma to the process. Each stimulation site was identified manually and the direction of a process movement was calculated relative to the center of the site. Similarly, process movements relative to the nearest neuron were calculated according to the center of the soma. Lastly, Directionality- and Transitional-indices (D- and T-indices) were calculated for each cell (Eq. 1 and 2) [40, 43, 44, 54, 99, 106, 107] separately with the length and position of processes manually drawn in ImageJ and processed in MATLAB (Figure 3b).

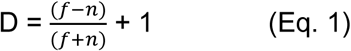

where *f* = # of processes away from electrode, n = # of processes near electrode.

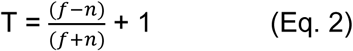

where *f* = length of longest process away from electrode, n = length of longest process near electrode.

In order to assess the directionality of process movements for a single microglia cell, a histogram of the angles of the process movements relative to the electrode or nearest neuron was plotted. This histogram was then fit with a probability distribution function using an Epanechnikov kernel function.

From there, the area under the curve (AUC) of the probability distribution function was calculated, with the area from 0 to 90 counting as toward the electrode, and 90 to 180 counting as away from the target. The polarity-index (P-index) was then calculated as a proportion of movements toward and away from the target (Eq. 3) and ranges from -1 to 1, where -1 indicates a microglia only extending processes away from the target and 1 represents a microglia only moving processes toward the target.

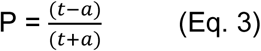

where *t* = AUC of movements toward target, *a* = AUC of movements away from target.

#### 2.5.2 Neural activity analysis

For each stimulation session, the average of each motion-corrected 1060 nm Z-stack was used to manually identify neurons and their profiles outlined using the ROI manager in ImageJ [73]. These ROIs and full ZT-series were subsequently imported into MATLAB for further fluorescence analysis. The fluorescence intensity of the neurons was filtered by taking the maximum intensity in each Z-stack to ensure that signals from somas not included in the full 20 um stack would not be adversely represented. The resulting fluorescent signal (F) had a sample for every 2.16 min representing the maximal neuronal calcium activity. To correct for noticeable photobleaching due to the length of the imaging session (100 min), the fluorescent signal for the full FOV of each imaging session was fit with a linear model excluding any stimulation timepoints. Each neuron’s fluorescent signal was then adjusted according to the linear model. Beyond the signal preprocessing, each neuron’s distance from the stimulation site was calculated for further spatiotemporal analysis.

Once the neuron signal has been preprocessed, the neuron was classified according to its activation and the subsequent temporal profile. Given that microglia participate in neuronal excitability through fractalkine and purinergic signaling [15, 61, 72, 108], identifying the behavior of a neuron provides more specific insight into how microglia may modulate its excitability. A neuron was considered activated if the ΔF/F0 was greater than the baseline F0 plus three times the standard deviation of the baseline period. Subsequent profile analyses determined 4 distinct profiles of activated neurons: depressed, baseline adapting, adapting, and non-adapting. Neurons were identified as depressed post-activation if the average calcium activity was less than -0.5 times the threshold identified earlier. Neurons were classified as baseline adapting if the average of the signal during stimulation was within one standard deviation of the baseline. Neurons were adapting if the average of the signal was between 1 and 3 standard deviations of the activation threshold. Finally, non-adapting neurons were identified if more than half of the stimulation timepoints exceeded the activation threshold, suggesting these neurons were active for more than 30 minutes. These classification criteria were mutually exclusive. Sample waveforms for these activation profiles are shown in Figure 6b.

**Figure 6.**
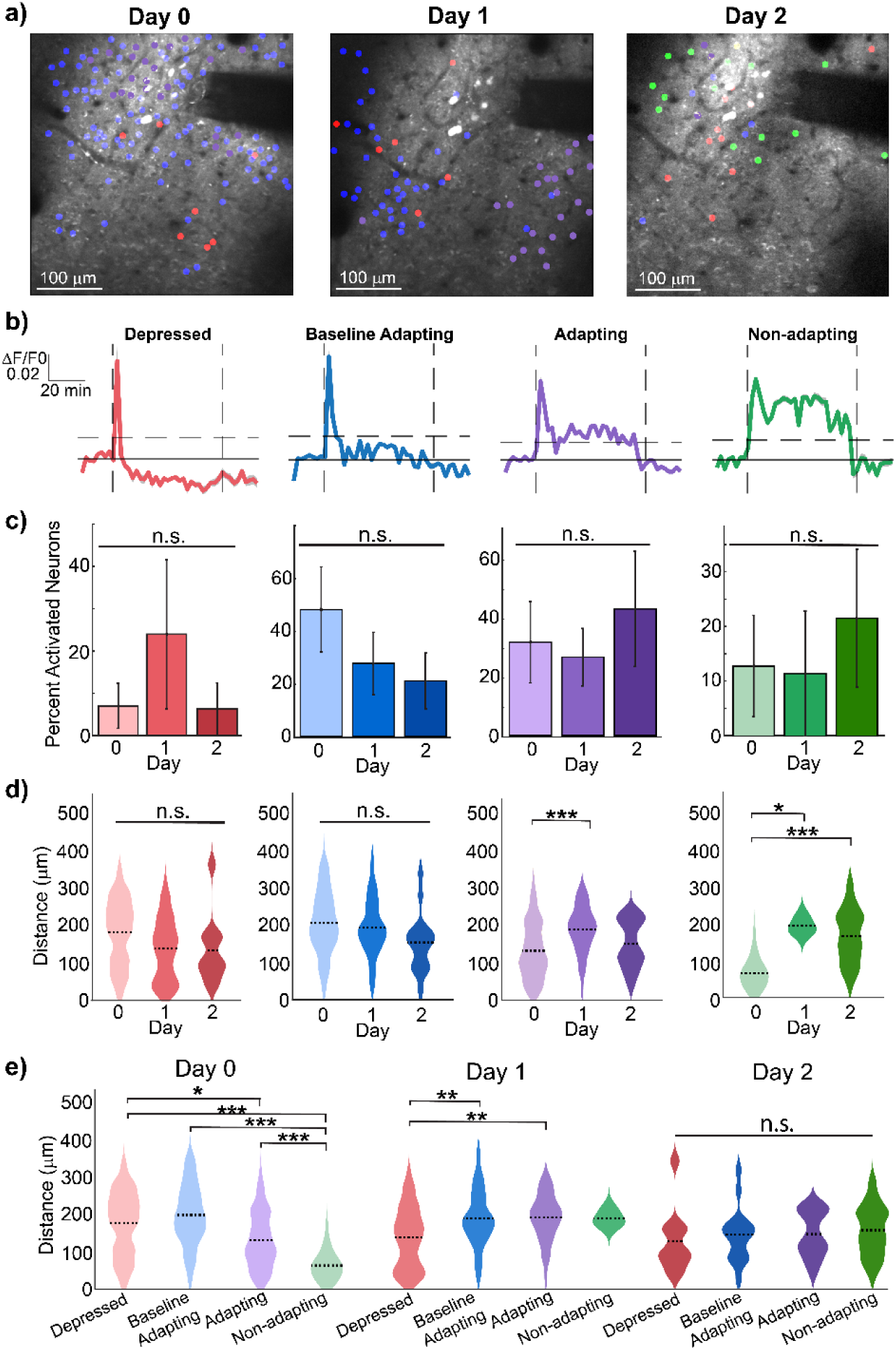
Neurons demonstrate spatially-dependent activation profiles during sustained 10-Hz ICMS. a) Representative chronic images of activated neuron profiles for spatial context. b) Representative calcium traces of differentiated neuron profiles, including stimulation onset (1^st^ vertical dashed line), offset (last vertical dashed line), and activation threshold (horizontal dashed line). c) The percentages of identified neuron profiles do not significantly vary across days (N = 4 mice, p = .543, .087, .742, .730). d) The neuron profiles exhibit less adaptation during the 1-h ICMS period are located further from the electrode across 2 days (N = 4 mice, Depressed n = 29, 39, 10, Baseline Adapting n = 191, 55, 15, Adapting n = 88, 38, 15, Non-adapting n = 27, 3, 31, One-Way ANOVA, Depressed p = .114, .294, .986, Baseline Adapting p = .601, .050, .225, Adapting p = 1.07e-4, .637, .170, Non-adapting p = .015, 1.41e-6, .814). e) On Day 0, less adapting neurons are found closer to the electrode (Kruskal-Wallis, p = .033, 3.62e-7, 1.22e-10, 8.23e-17, 4.13e-4). By Day 1, depressed adapting neuron profiles move closer to the electrode while adapting neuron profiles move further away (p = .004, .008, .652, 1, 1, 1). On Day 2, there is no significant difference in the spatial distribution of the neuron profiles (p = .931, .919, .682, 1, .957, .967).

Non-activated neurons also exhibited distinct temporal profiles, with a subset showing immediate depressed activity at the onset of stimulation. These depressed non-activated neurons (Supp. Figure 6b) were defined as having average calcium activity less than −0.5 times the activation threshold, without ever exceeding the activation criterion. This contrasts with depressed post-activation neurons, which first crossed the activation threshold before subsequently falling below baseline activity.

While neurons exhibited distinct activation profiles during stimulation, there were also two distinct profiles of neuronal calcium activity post-stimulation (Figure 8a). Neurons were classified as depressed post-stimulation if the average calcium activity was less than -0.5 times the activation threshold.

Otherwise, neurons were classified as baseline post-stimulation when the activity was within 0.5 standard deviations of the baseline activity.

### 2.5.3 Microglia-neuron interaction analysis

To examine microglia-neuron interactions during stimulation, microglia were overlaid with the neuronal ROIs identified earlier and each interaction was manually recorded alongside the timing of the microglia process movement. Interactions were classified into connections and disconnections whenever microglia processes touched a neuron soma or retracted away from the soma. It is crucial to note that any disconnections would require the microglia to already be in contact with the neuronal soma.

#### 2.5.4 Statistics

All data in violin plots ([109] Violin plots for Matlab) show the mean (horizontal line). All statistical analyses were performed in MATLAB with an alpha level of 0.05 and Bonferroni post-hoc correction for multiple comparisons. Prior to any analysis of variance (ANOVA), the assumption of homogeneity of variance was tested via Levene’s Absolute Test, and if significant, a non-parametric alternative like the Kruskal-Wallis test was employed. Kolmogorov-Smirnov tests were used in the case of distributional and angular analyses to identify differences in behavior. Error bars are standard error unless otherwise stated.

## Results

Two-photon imaging in Cx3cr1^GFP^/jRGECO1a mice enabled simultaneous tracking of neuronal calcium activity and microglial dynamics during prolonged 10-Hz ICMS (1 h). Four distinct neuronal adaptation profiles were identified: (a) *non-adapting* neurons maintaining suprathreshold activity throughout 60-min stimulation, (b) *adapting* neurons showing progressive activity reduction but remaining above baseline, (c) *baseline-adapting* neurons returning to pre-stimulation activity levels during stimulation, and *depressed* neurons exhibiting sub-baseline activity post-activation. Microglial processes dynamically responded to ICMS, extending toward stimulation sites and active neurons, with microglia-neuron contact frequency inversely correlating with neuronal adaptation magnitude. Mechanistic insights were further gained by comparing acute (3 h) and chronic (24–48 h) post-implantation responses, which revealed that microglial encapsulation of electrodes coincided with neuronal hypoactivity. Using cellular-resolution imaging and biphasic stimulation (2 nC/phase; within established safety thresholds [110]), we demonstrate that microglia may modulate ICMS-induced network activity. These findings challenge neuron-centric paradigms and support a model in which glia function as computational elements influencing stimulation efficacy.

### 3.1 Chronic microelectrode insertion drives and directs faster microglial process movements

Microglia respond to injuries in the brain, including those caused by lesions, stroke, or microelectrode insertion. Unlike transient injuries, microelectrode implants represent a persistent foreign body that continuously interacts with the surrounding neural tissue, leading to prolonged glial responses such as encapsulation via sheath formation [55, 111, 112]. Within 3 h of electrode insertion, microglia extend their processes toward the implant [54, 113], initiating a cascade that includes phagocytosis of damaged cells and clearance of injury-related debris. To isolate the contribution of microglia to the neuronal response during ICMS, it was necessary to first characterize microglial behavior following chronic electrode implantation in the absence of stimulation (Figure 1C-D). This baseline assessment enabled evaluation of microglia-neuron interactions specific to ICMS rather than injury alone.

Interestingly, microglia surveillance, assessed by the ratio of measured extensions to retractions remained consistent over days (Figure 1C). However, there were significantly more extensions than retractions on Day 1 (Supp. Figure 1B; p = 0.038). This corresponds with the peak of microglial process migration commonly observed in the first day post-injury [55, 114–117]. Notably, the ratio of extensions to retractions remained unchanged and the total number of process movements per microglia did not increase during the post-implantation period (Supp. Figure 1A). Additionally, this ratio was independent of the distance of microglia from the electrode or the time post-implantation (Supp. Figure 1D). These results suggest that microglial surveillance, in terms of the balance of process movements, does not shift despite the established injury response and the ongoing chronic injury caused by the microelectrode.

Instead, microglia behavior appears to change in response to electrode injury, specifically in terms of the speed of process movements. Both process extensions and retractions exhibit increased movement speeds post-implantation (Figure 1D extension speeds: Day 0: 0.733 ± 0.0138 µm/min, Day 1: 0.815 ± 0.0130 µm/min, Day 2: 0.888 ± 0.0140 µm/min, retraction speeds: Day 0: 0.778 ± 0.0160 µm/min, Day 1: 0.941 ± 0.0242 µm/min, Day 2: 0.933 ± 0.0174 µm/min). However, the speed of these process movements is inversely proportional to the number of movements, with process retractions consistently being faster than extensions (Supp. Figure 1C). Furthermore, like the balance of extensions and retractions, the speed of both extensions and retractions does not vary with the distance of microglia from the electrode insertion site (Supp. Figure 1E).

To better understand how microelectrode insertion affects microglia behavior beyond standard metrics of process movement, we assessed the directionality of process movements relative to the electrode, particularly the stimulation site (Figure 2A). Distinct behavioral changes were observed based on the distance of microglia from the electrode and directionality of process extensions relative to the electrode were quantified. Microglia located within 150 µm of the electrode extend more frequently towards the electrode and site of injury compared to microglia further away (Figure 2B). Only extensions were analyzed in this case, as retractions were typically in the opposite direction of extensions, providing little additional information on microglial behavior (Supp. Figure 2A-B).

In addition to analyzing the distribution of microglia extension angle relative to the electrode, we established a condensed metric – the microglial process polarity index – to indicate whether the average microglia extends toward (positive) or away from (negative) the electrode (Eq. 3, Supp. Fig 2C-D). The polarity index was consistent with the distribution of extension angles (Figure 2C, n = 18, 25, and 6 microglia, polarity index: Near: 0.216 ± 0.0643, Mid: -0.053 ± 0.0647, Far: -0.224 ± 0.1879), indicating that microglia within 150 µm of the electrode extend toward the stimulation site more frequently than microglia further away. Since these extensions occurred during baseline, the more common extensions toward the electrode are most likely a function of the injury caused by insertion.

Interestingly, the distribution of these extensions toward the electrode peaked on Day 1 for microglia within 150 µm of the stimulation site (Figure 2D), although the microglia did not appear polarized in terms of the number of extensions toward the electrode (Figure 2E, n = 18, 10, and 11 microglia, polarity index: Day 0: 0.216 ± 0.064, Day 1: 0.451 ± 0.1134, Day 2: 0.321 ± 0.0977). These results suggest that the distribution of microglia extensions toward the electrode are influenced by the injury response, though additional factors, such as surrounding tissue and cellular behavior, may also play a role in modulating microglia process extensions.

### 3.2 Prolonged low frequency ICMS does not affect microglia morphology, but modulates and directs process extensions

Previous studies have suggested that low-frequency electrical stimulation (Figure 3A), had ameliorating effects on microglial activation post-stimulation, potentially reducing the negative consequences of inflammation [15, 16, 118]. However, it remains unclear whether stimulation alters microglia inflammatory phenotypes during ongoing stimulation. Microglia morphology is a direct indicator of their activation state, with homeostatic and resting microglia presenting as ramified with substantial branching, while inflamed microglia adopt a more amoeboid morphology with fewer processes [17, 54, 119, 120]. Therefore, we next quantified whether microglia morphology changed during or following stimulation. Directionality- and transitional-indices (D- and T-indices) have been used to quantify microglial morphology in response to microelectrode insertion [44] (Figure 3B). In this study, no significant change in the D-index was observed following stimulation across all days (Figure 3C), nor were there any significant changes in the T-index over the same period (Figure 3D). Thus, the 1h application of low-frequency ICMS did not significantly alter the morphology of surveilling microglia during or after stimulation.

Interestingly, microglia morphology exhibited some significant dependence on distance on Day 1, with lower D- and T-indices observed for microglia closer to the electrode (Supp. Figure 3B, 3D; D-Index: p = 0.046; T-Index: 150-300: p = 0.0034,>300: p = 0.0007). This distance-dependence pattern aligns with previous studies suggesting that microglia near the electrode undergo morphological changes in response to the damage caused by microelectrode insertion [54]. Despite these distance-related differences, a significant decrease in T-index was observed on Day 1 for microglia located more than 300 µm away from the electrode, occurring between the onset of stimulation and 20 min post-stimulation (Supp Figure 3F; p = 0.042). These mixed results suggest that stimulation may have a modulatory effect on microglia activation, but there is no consistent effect of low-frequency ICMS on microglia morphology or activation.

Since ICMS did not seem to affect microglia activation and morphology, we next examined whether stimulation influenced microglia behavior in terms of process movement speed. Although there were subtle fluctuations in process speeds throughout stimulation, extension speeds were only significantly modulated on Day 2, where a decrease was observed after the offset of stimulation (Figure 3E; Early v Post: p=0.038 and Late v Post: p=0.036). In contrast, retraction speeds were significantly modulated on Day 0 where they increased following the offset of ICMS (Figure 3F; p = 0.042).

Given that stimulation demonstrated some effects on microglia process behavior, we further investigated whether these behavioral changes might be related to microglia extensions in relation to the site of stimulation. On Day 0, the onset of stimulation increased the distribution of microglia extensions toward the electrode, but did not polarize the microglia extensions toward the electrode (Figure 4A). For both Day 1 and 2, microglial extensions were already predominantly directed toward the electrode site, and no shift in distribution or polarization occurred due to stimulation (Figure 4B-C). Similarly, microglial retractions largely opposed the direction of extensions and exhibited no significant directional shifts (Supp. Figure 4A-C). The observed changes in extension direction suggest that microglia are likely responding to the electrical stimulation, but there is no definitive evidence that ICMS is driving the extensions toward the electrode alone. Neurons may communicate with microglia through chemical signaling, such as fractalkine, and since more neurons are activated closer to the electrode, neuronal activity could also contribute to microglial behavior changes.

### 3.3 Microglia process extensions migrate toward neuronal activation during low frequency ICMS

Given the observed shifts in microglia process behavior in response to ICMS, we next explored how these behaviors are linked to neuronal activity. Fewer neurons were identified as activated over time, with a significant decrease (p = 0.0125) between Day 0 and 2 post-implantation (Figure 5C, Day 0: 71.93 ± 7.81%, Day 1: 38.10 ± 9.90%, Day 2: 23.29 ± 9.61%). The distance of activated neurons from the electrode site was significantly lower on Day 2 than Day 1 (Figure 5D, Day 0: 173.41 ± 4.98 µm, Day 1:179.14 ± 6.66 µm, Day 2: 152.96 ± 8.45 µm), which aligns with previous findings suggesting that the inflammation around a microelectrode insertion decreases the response to stimulation [84, 112, 121, 122]. To assess microglia responses to neuronal activity, we defined microglia processes as being near neurons if they were within 20 µm, thus capable of interacting with the neurons during any of the 20-min periods of stimulation (Figure 5E) given that microglia were calculated to have a maximum process speed of 0.941 ± 0.0242 µm/min (Figure 1D). Interestingly, microglia process extensions responded inversely to their extensions towards the electrode, with significantly more extensions toward activated neurons during stimulation on Days 1 and 2, although the microglia behavior does not appear more significantly polarized (Figure 5G; extension p =.021, polarity index p = 0.149; Figure 5H; extension p = 0.002, polarity index p = 0.313). As before, microglia retractions tended to oppose extensions toward activated neurons, showing no significant variations in retraction direction relative to activated neurons (Supp. Figure 5). These results suggest that microglia are more responsive to neuronal activation in the surrounding tissue after the acute phase of their injury response, rather than to the electrical stimulation itself.

### 3.4 Neuronal activity decreases with increasing microglia-neuron interactions

Building on the observed changes in microglial behavior and their potential interaction with neuronal activity, we next assessed how neuronal activation profiles vary in relation to the electrode and microglial response. Neurons exhibit different activity profiles during ICMS depending on distance from the electrode [9, 11–13, 73, 74, 123]. A sample of chronic spatial neuron activity revealed a varied and seemingly inconsistent distribution of neuronal activation profiles (Figure 6A). Four distinct activation profiles emerged when examining the neurons: 1) depressed neurons, which show diminished calcium activity post-activation, 2) baseline adapting neurons, which adapt back to baseline levels during stimulation 3) adapting neurons, which exhibited decreased calcium activity but still above a threshold of one standard deviation, and 4) non-adapting neurons, which demonstrated sustained activation throughout stimulation (Figure 6B). Despite the spatial variability in activation profiles, there were no significant or consistent changes in the proportion of these profiles across the days during low-frequency ICMS (Figure 6C, in order Day 0, Day 1, Day 2: Depressed: 7.17 ± 5.486%, 24.72 ± 18.234%, 6.41 ± 6.41%, Baseline Adapting: 47.88 ± 16.049%, 27.57 ± 11.681%, 20.97 ± 10.602%, Adapting: 32.96 ± 14.355%, 27.64 ± 10.098%, 44.67 ± 20.337%, Non-adapting: 14.35 ± 8.660%, 10.10 ± 10.714%, 20.34 ± 11.778%). However, despite the stable proportions, non-adapting neurons were located significantly farther from the electrode (Figure 6D, in order Day 0, Day 1, Day 2 Depressed: 182.39 ± 16.53 µm, 139.09 ± 13.52 µm, 134.20 ± 29.48 µm, Baseline Adapting: 204.82 ± 6.11 µm, 192.64 ± 10.48 µm, 152.89 ± 20.44 µm, Adapting: 135.495 ± 8.26 µm, 194.11 ± 10.50 µm, 153.99 ± 16.99 µm, Non-adapting: 65.25 ± 7.66 µm, 191.76 ± 12.87 µm, 164.99 ± 13.12 µm). Both adapting (p = 0.0002) and non-adapting (p=0.015) neurons shifted significantly away from the electrode by Day 1, which opposes the typical migration of microglia towards an injury site (the electrode). Interestingly, on Day 0, non-adapting neurons were localized significantly closer to the electrode compared to other profiles (Figure 6E; p < 3.62e-7), and adapting neurons were also closer to the electrode than depressed neurons (Figure 6E; p = 0.033). By Day 1, depressed neurons were located significantly closer to the electrode than baseline (p=0.004) or adapting neurons (p=0.008; Figure 6E), suggesting that the spatial distribution of sustained neuronal activation may shift as part of the chronic injury response. Although neuron recruitment and magnitude of activation is often related to distance from the electrode, the proportion and spatial localization of depressed neurons do not exhibit significant changes across days (Supp. Figure 6). While neuron activation seems to be influenced by proximity to the electrode, the lack of change in proportions and distances for depressed neurons, coupled with variability across animals, indicates that additional factors are at play. Microglia-neuron interactions are the most likely candidates for influencing neuronal activation in this context.

To further investigate the role of microglia in modulating neuronal activity during ICMS, we examined the spatial and temporal dynamics of microglia-neuron interactions. By isolating neuron somas and identifying when microglia processes made and broke contact with nearby neurons (Figure 7A), we quantified both the timing and frequency of microglia-neuron interactions. Across all three days, non-activated neuron ROIs were more prevalent (Figure 7B) prompting an initial comparison between activated and non-activated neurons. This analysis revealed no significant differences in the timing of microglia-neuron interactions between the two groups (Figure 7C).

**Figure 7.**
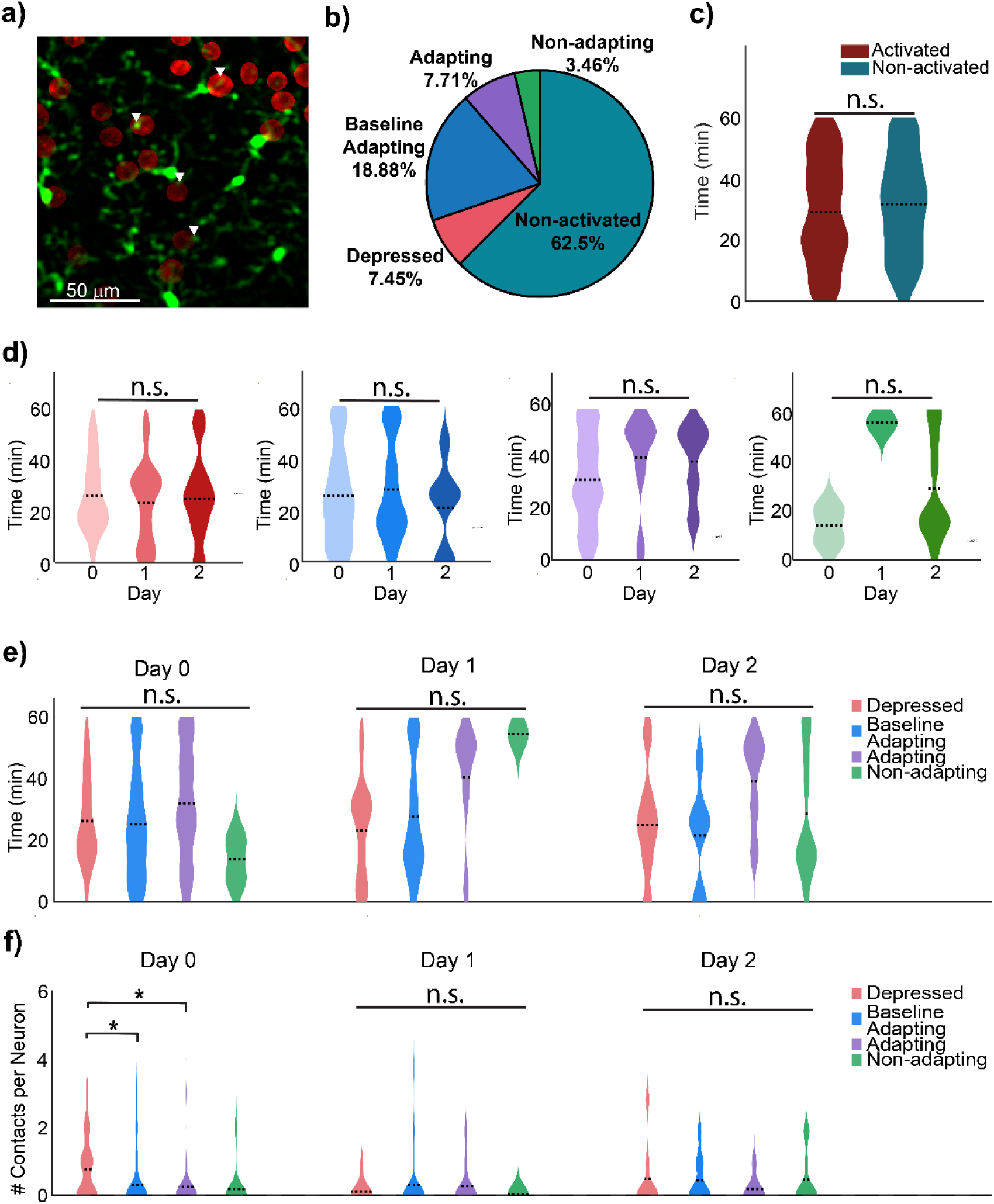
Microglia contact the somas of depressed activated neurons more frequently than other profiles on Day 0. a) Two-photon microscopy of microglia in proximity to neuronal somas was used to measure microglia-neuron interactions. b) Percentage of neuron profiles across all chronic timepoints. c) Peri-stimulation analysis of the timing of microglia interactions with activated and non-activated neurons, showing no significant difference between the timing of interactions (N = 99, 123 neurons, n = 45, 83 peri-stimulation interactions, One-Way ANOVA, p = 0.422). d) Peri-stimulation timing of microglia-neuron interactions with identified neuron activation profiles, demonstrating no significant changes in when microglia interact with specific neuron profiles over days (Depressed N = 13, 8, 7 neurons, n = 16, 10, 6, peri-stimulation interactions, Baseline Adapting N = 51, 15, 5 neurons, n = 69, 20, 6, peri-stimulation interactions, Adapting N = 18, 8, 3 neurons, n = 19, 12, 5, peri-stimulation interactions, Non-adapting N = 1, 1, 11 neurons, n = 2, 1, 10, peri-stimulation interactions, One-Way ANOVA, p = 0.901, 0.501, 0.395, 0.287). e) Peri-stimulation timing of microglia interactions with different neuron activation profiles is not significantly different (One-Way ANOVA, p = 0.379, 0.074, 0.434). f) Microglia interact more frequently with neurons that demonstrate more suppressed activation profiles on Day 0 (Depressed N = 20, 20, 8, n = 16, 10, 6, peri-stimulation interactions, Baseline Adapting n = 124, 36, 9 neurons, Adapting n = 56, 22, 11neurons, Non-adapting n = 11, 1, 21 neurons, One-Way ANOVA, Day 0 p = 0.046, 0.044, Day 1 p = 0.726, Day 2 p = 0.747).

**Figure 8.**
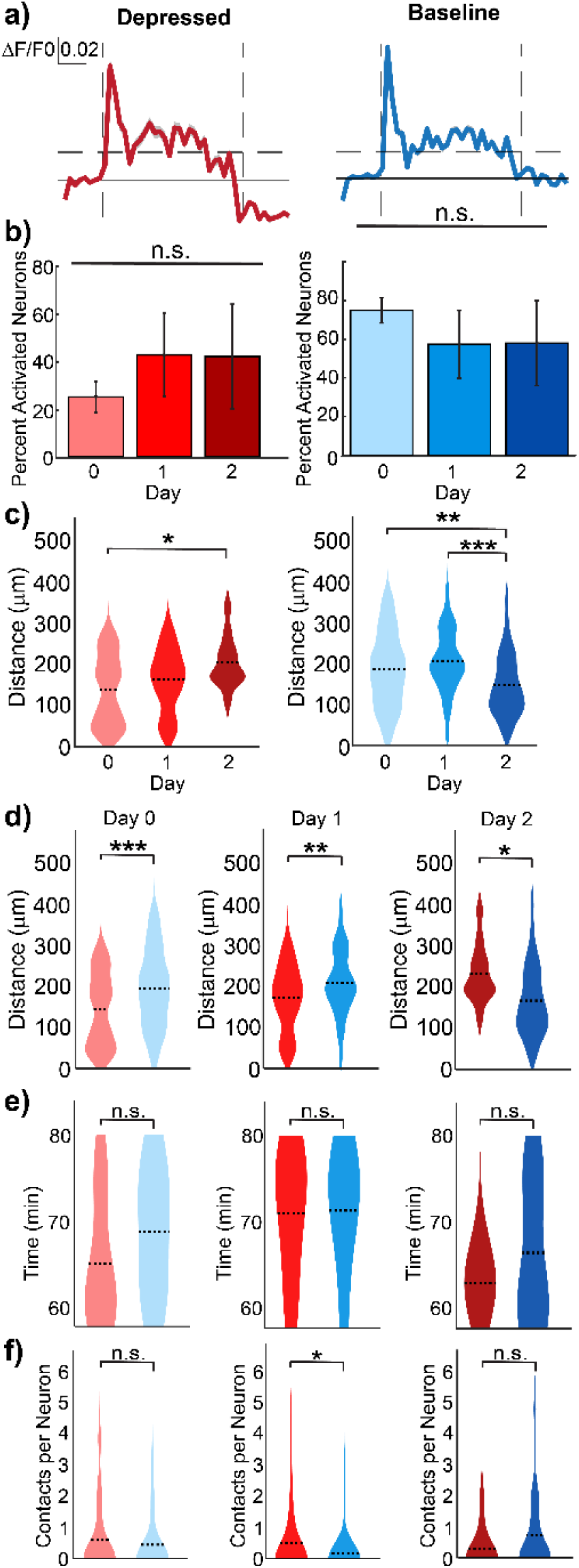
Microglia may regulate neuronal activity post-ICMS through contact with neuronal somas. a) Representative calcium traces of neurons with baseline or depressed activity profiles following stimulation. b) The proportion neurons exhibiting baseline versus depressed post-stimulation profiles did not significantly change across the days (N = 4 mice, One-Way ANOVA, p = 0.708). c) Neurons with depressed post-stimulation activity shifted farther from the electrode across days (n = 72, 80, 11 depressed neurons, One-Way ANOVA, p = 0.027) while neurons with baseline post-stimulation profiles shifted toward the electrode (n = neurons, One-Way ANOVA, p = n = 263, 60, 72 baseline neurons, One-Way ANOVA, p = 0.0025, 0.0004). d) On Days 0 and 1, depressed post-stimulation neurons were significantly closer to the electrode compared to baseline neurons (One-Way ANOVA, p = 0.0001, 0.002), but by Day 2, baseline neurons were significantly closer (One-Way ANOVA, p = 0.018). e) The timing of microglia-neuron interactions post-stimulation did not significantly differ by activation profile on any day (Depressed N = 53, 51, 10 neurons, n = 10, 6, 1 poststimulation interactions, Baseline N = 158, 31, 46 neurons, n = 32, 6, 10 poststimulation interactions, One-Way ANOVA, p = 0.157, 0.935, 0.689). f) On Day 1, microglia contacted depressed post-stimulation neurons significantly more frequently than baseline neurons (Kruskal-Wallis, p = 0.040). No significant differences were observed on Days 0 or 2 (One-Way ANOVA, p = 0.3266, 0.3026). No significant differences were observed on Days 0 or 2 (One-Way ANOVA, p = 0.327, 0.303).

Furthermore, the timing of interactions did not significantly change across days for any of the activation profiles (Figure 7D), nor did it differ between activation profiles at individual time points (Figure 7E). Although some differences in timing were visually apparent, the small number of adapting and non-adapting neurons may have limited statistical power.

Given that interaction did not appear to influence neuronal activation profiles, we next considered whether microglia-neuron interaction frequency during stimulation might reflect a modulatory or homeostatic role for microglia, as proposed in prior studies [61, 92–94]. On Day 0, microglia interacted significantly more frequently with depressed neurons than with baseline (p=0.046) and adapting (p = 0.044) (Figure 7F, Day 0: depressed: 0.75 ± 0.204, baseline: 0.29 ± 0.060, adapting: 0.25 ± 0.109, and non-adapting: 0.18 ± 0.182 contacts per neuron, suggesting a potential suppressive effect on neuronal activity. However, when considering that non-activated neurons were contacted even more frequently than depressed neurons (Supp. Figure 7C), the data are more consistent with a microglial role in maintaining homeostasis in response to stimulation.

Another critical factor of ICMS is neuronal activity following stimulation. At lower frequencies and shorter durations (1-30 s), a larger proportion of neurons typically return to their baseline activity [11, 73, 74]. Alternatively, some neurons exhibit a significantly depressed calcium signal post-stimulation. Both response types were observed (Figure 8A), and the proportions of baseline and depressed post-stimulation profiles did not significantly change across days (Figure 8B). However, by Day 2, a spatial inversion emerged. Depressed post-stimulation neurons were located further from the electrode, whereas baseline post-stimulation neurons were closer (Figure 8C in order Day 0, Day 1, Day 2 Depressed: 137.44 ± 9.98 µm, 161.55 ± 8.59 µm, 203.90 ± 18.97 µm, Baseline: 183.27 ± 5.58 µm, 202.60 ± 9.79 µm, 145.18 ± 8.99 µm). This trend was confirmed by direct comparisons, where depressed neurons were significantly closer to the electrode on Days 0 and 1, but significantly farther on Day 2 (Figure 8D; Day 0: p = 0.0001; Day 1: p = 0.002; Day 2: p = 0.018). In contrast to the spatial changes, the timing of microglia contacts with neuronal somas post-stimulation did not differ significantly across days (Figure 8E). Interestingly, microglia interacted significantly more frequently with depressed post-stimulation neurons on Day 1 (Figure 8F; p = 0.040; in order Day 0, Day 1, Day 2 Depressed: 0.60 ± 0.156, 0.49 ± 0.149, 0.30 ± 0.213, Baseline: 0.44 ± 0.079, 0.16 ± 0.115, 0.74 ± 0.190 contacts per neuron). These data suggest that microglia may help restore homeostasis by reducing excitability in neurons that experienced prolonged activation [75, 124].

## Discussion

ICMS evokes heterogeneous neuronal responses that evolve over time, reflecting both immediate circuit-level effects and dynamic neuromodulation. This study demonstrates that microglia-neuron interactions are temporally and spatially regulated during and after stimulation, with microglia preferentially contacting neurons exhibiting suppressed or altered activation states. Using *in vivo* two-photon imaging over three consecutive days, we tracked microglial process contacts with neuronal somas and categorized neuronal calcium responses into distinct activation profiles during 10-Hz ICMS. Analyses revealed that while the *timing* of microglia-neuron interactions did not correlate with activation states, the *rate* of interactions was significantly elevated for suppressed neurons during and after stimulation. These contact rates were particularly enriched in activated neurons classified as "depressed" based on ΔF/F reductions and spatially localized outside the immediate vicinity of the electrode (Fig. 6D–E), supporting the hypothesis that microglia prioritize metabolically or synaptically compromised targets. These interactions may be exerting a homeostatic influence rather than merely suppressing activity, suggesting a more nuanced role for microglia in shaping cortical network responses to electrical stimulation. Together, these findings highlight the importance of considering glial dynamics in the interpretation and optimization of ICMS-based neuromodulation strategies.

### 4.1 Low frequency ICMS does not induce morphological microglia activation

Our study demonstrates that prolonged one-hour 10-Hz ICMS does not induce classical microglial activation, as evidenced by the absence of morphological changes in directionality- and transition-indices (D- and T-indices; Fig. 3). Instead, microglial processes dynamically responded to ICMS by extending toward both the stimulating electrode and active neurons (Fig. 4 and Fig. 5F), with contact frequency inversely correlating with neuronal adaptation magnitude (e.g., depressed neurons exhibited more microglial interactions; Fig. 7F). This targeting was selective despite no changes in average D- or T-indices (Fig. 3C–D), suggesting that microglia adapt their motility to local neurochemical cues without undergoing global activation. Notably, this behavior was associated with neuronal activity, suggesting that microglial responses are primarily driven by chemical signaling (e.g., ATP, fractalkine) released by neurons, rather than the electric field directly [15, 61, 72, 108]. This is consistent with established roles for P2Y12 and CX3CR1 receptors in guiding microglial surveillance in response to ATP and fractalkine, respectively, likely mediating the preferential process extension toward activated neurons observed on Days 1 and 2 (Fig. 5F–H). Although transient shifts in process speed were detected (e.g., faster retractions on Day 0, slower extensions on Day 2; Fig. 3), these effects were inconsistent across days, further supporting the idea that 10-Hz ICMS alone is insufficient to drive sustained microglial reactivity. Microglia-neuron interactions appeared homeostatic as neurons with depressed calcium activity post-stimulation were contacted more frequently during stimulation (Fig. 7), implying a role in metabolic recovery or synaptic tuning rather than simple inhibition. This interpretation is further supported by the fact that less suppressed neurons gradually appear further away from the electrode, implying an active compensatory or monitoring role for microglia.

These findings challenge the assumption that electrical stimulation could directly activate microglia and instead highlight their role as adaptive regulators of neuronal networks. Microglia seem to facilitate a balance in neuronal activity, rather than simply inducing neuroinflammation. However, these interactions may be accompanied by subtle transcriptomic remodeling, such as upregulation of anti-inflammatory markers (e.g., Arg1, IL-10) or synapse-modulating factors (e.g., TREM2, IGF1), which were not assessed in this study but should be explored using post hoc RNA-seq or immunostaining of targeted cells. The subtlety of the response also underscores the importance of stimulation frequency in modulating glial engagement. While 10-Hz ICMS did not provoke morphological changes, higher frequencies (e.g., 40–100 Hz) may drive stronger neuronal adaptation (Fig. 6B) and surpass a threshold for microglial reactivity [11, 13, 73]. Parametric variation of frequency and duty cycle could identify thresholds for transitioning between surveillance, homeostatic modulation, and inflammatory states. Such profiles would inform precision programming of ICMS paradigms that minimize adverse glial responses.

Given that microglial encapsulation of electrodes coincides with progressive neuronal hypoactivity in chronic implants [84, 113], prolonged 10-Hz ICMS over days or weeks could reveal delayed glial adaptations, such as process retraction, phagocytic engagement, or altered cytokine release. These adaptations may underlie the gradual decline in stimulation efficacy observed in long-term neuroprosthetic applications [40]. Extending longitudinal two-photon imaging beyond three days (e.g., to Day 7 or Day 14) would determine whether transient process accelerations (Fig. 3E–F) evolve into structural reorganization, such as sheath formation or synaptic pruning.

Future experiments should focus on tracking microglial dynamics in long-term neuroprosthetic applications, paired with high-resolution imaging of sheath formation around electrodes. Such studies will help determine whether low-frequency ICMS accelerates or mitigates inflammatory encapsulation. Additionally, combining chronic ICMS with longitudinal electrophysiology could clarify whether microglial interactions predict late-phase neuronal silencing or aberrant plasticity. For example, spatial remapping of depressed neuron distributions post-stimulation (Fig. 8C–D) may correlate with local field potential changes and could be modulated by prior microglial engagement. Integration of ΔF/F imaging with field recordings would offer a multi-modal metric of microglia-neuron feedback. Such studies would bridge the gap between acute microglial surveillance and chronic circuit modulation, informing strategies to sustain stimulation efficacy through glial-targeted interventions.

While 10-Hz ICMS does not overtly activate microglia, it engages them as subtle modulators of neuronal adaptation. Future work should prioritize parametric studies of stimulation paradigms, and chronic applications to unravel glial contributions to neuromodulation efficacy. Refining stimulation protocols to explicitly account for microglial roles may enable more stable and precise control of neural circuits.

### 4.2 Low frequency ICMS modulates microglia behavior

Although a one-hour, 10-Hz ICMS does not elicit classical morphological activation of microglia (no change in D- or T-indices; Fig. 3C–D), it does subtly alter microglial dynamics in ways that depend on local circuit activity (Fig. 5). During prolonged 10-Hz stimulation, microglial processes exhibit directional plasticity, extending more frequently toward the electrode shank and along trajectories that mirror regions of heightened neuronal calcium activity (Fig. 5F–H). We observed that microglial processes actively extended toward both the stimulation site and activated neurons during prolonged 10-Hz ICMS, with contact frequency inversely correlating with neuronal adaptation magnitude (Fig. 7). Critically, these process extensions occurred without overt changes, indicating that low-frequency ICMS engages microglia primarily by influencing their surveillance patterns rather than driving a full activation phenotype[15, 61, 72, 108]. These findings challenge neuron-centric models of neuromodulation and position microglia as adaptive partners in shaping ICMS outcomes.

Quantitative analysis of microglial motility revealed that, on Day 2 post-implantation, average process extension speed decreased slightly after 10-Hz stimulation (Day 2; Fig. 3E), while retraction speed increased after stimulation on Day 0 (Fig. 3F). These modest, time-dependent shifts in process kinetics suggest that low-frequency ICMS can alter microglial surveillance behavior, potentially via ATP/adenosine signaling cascades rather than direct electrical depolarization [61, 92–94]. For example, field-evoked glutamate release and downstream ATP hydrolysis to adenosine are known to regulate P2Y12R–A1R pathways, dampening microglial ramification without triggering a full-scale inflammatory profile[125, 126].

Even in the absence of morphological activation, microglia in proximity (<150 µm) to the stimulating electrode exhibited significant polarization toward the implant during post-implantation “baseline” periods (Fig. 2C–E). By Day 1, the polarity index for extensions peaked near the electrode, reflecting an injury response (Fig. 2D–E). Notably, 10-Hz stimulation did not significantly amplify overall polarization but instead modulated the balance of process extension and retraction in a directionally structured manner (Fig. 5). As neurons near the electrode exhibited depressed activity over time (Fig. 6D–E), microglial processes shifted away from these regions, suggesting a possible link between sustained low neuronal activity and reduced microglial engagement. On Day 1, microglia showed increased engagement with neurons exhibiting depressed post-stimulation ΔF/F (Fig. 8F), implying that microglia may sense and respond to regions of metabolic stress. This aligns with the shift in neuronal activity profiles (Fig. 7), where depressed neurons are more frequently contacted by microglia post-stimulation (Fig. 7). The observed temporal pattern aligns with prior observations that, as neurons near an active electrode experience metabolic strain, they exhibit decreased BOLD-OIS amplitudes and eventual silencing [127]. Microglia likely detect local shifts in extracellular adenosine or other metabolic byproducts and extend processes to these regions of metabolic stress, microglia may aid in clearing extracellular metabolites, modulating extracellular ion balance, or releasing trophic factors to restore homeostasis, actions analogous to their roles in injury repair and synaptic refinement [18, 86]. As BOLD-OIS signals decrease near implants [127], microglial engagement may help stabilize network function without inducing pro-inflammatory phenotypes.

The progressive shift of depressed neurons away from the electrode (Fig. 6D), combined with microglial redistribution (Fig. 2D–E) suggests that low-frequency ICMS fosters a dynamic equilibrium: microglia monitor metabolic stress and help preserve tissue homeostasis without driving overt gliosis. This interpretation is supported by studies demonstrating that 10 Hz ICMS suppresses pericyte calcium activity in a frequency-dependent manner, suggesting coordinated neuromodulation of non-neuronal cells [128]. Such responses are unlikely to arise from direct depolarization, further supported by the fact that the direct activation volume appears to be roughly 10-20 µm from the electrode site as measured by iGluSnFR [129]. Instead, the collective findings implicate purinergic or adenosine-mediated signaling cascades across glial-vascular networks. This may mitigate the encapsulation and signal attenuation observed in chronic implant studies [84, 113]. Therefore, neuromodulation protocols should account for microglial dynamics by tailoring ICMS parameters (frequency, duty cycle, amplitude) to align with microglial surveillance windows to stabilize tissue responses and prolong functional integration.

Incorporating biomaterials engineered to optimize charge transfer and modulate microglial membrane potentials, such as conductive polymer coatings that limit ATP release, may further harness these glial mechanisms to support biocompatible long-term implants

### 4.3 Mhicroglia-neuron interactions

Our data indicate that, even at low-frequency (10 Hz) ICMS, microglia may serve as active intermediaries in cortical circuits, extending processes toward both the electrode and nearby active neurons (Fig. 5). Although this stimulation paradigm does not induce classical inflammatory activation in surveilling microglia (no change in D- or T-indices; Fig. 3C–D), microglial processes nevertheless exhibit directional plasticity, suggesting a role in fine-tuning synaptic function rather than merely reacting to injury.

Microglia express diverse neurotransmitter receptors and voltage-sensitive receptors [38, 130–132], which may enable direct sensitivity to ICMS-evoked glutamate or GABA. Neuronal activity under 10 Hz ICMS may lead to the release of several neurochemicals that likely guide microglial behavior. Elevated extracellular ATP released by hyperactive neurons engages microglial P2Y12 receptors, promoting process extension toward active synapses [125, 126]. ATP hydrolysis to adenosine via microglial CD39/CD73 triggers A1 receptor signaling on neurons, which dampens excitability and may provide feedback to modulate microglial surveillance [125, 133]. Likewise, fractalkine (CX3CL1) released from stimulated neurons binds microglial CX3CR1, regulating chemotaxis and synaptic pruning [22, 134, 135]. It is important to note that our animal model was heterozygous for CX3CR1, preserving one functional allele. Excess glutamate, abundant during sustained activation, can engage microglial mGluRs and, if unregulated, drive pro-inflammatory signaling [136, 137]. Together, these signals coordinate microglial process extension and retraction to maintain network stability.

As expected, microglial contact frequency is highest for neurons exhibiting the greatest adaptation. Depressed or highly adapting neurons receive more frequent microglial contacts (Fig. 7F; Fig. 8F). Physical microglia–neuron interactions during ICMS likely depend on local concentrations of ATP, adenosine, and fractalkine: ATP and fractalkine steer microglia toward regions of high firing, whereas adenosine may signal to restrain microglial over engagement once neuronal excitability subsides[125, 126, 138]. Additionally, low-frequency ICMS (2–20 Hz) is known to enhance BDNF and other trophic factor release, which suppresses microglial pro-inflammatory cytokines and promotes angiogenesis [49, 50, 95]. By contrast, higher frequencies (e.g., 40 Hz) induce robust microglial gene expression and morphological changes [139, 140]. Although we did not observe morphological activation at 10 Hz (Fig. 3C–D), this frequency-dependent balance between trophic and inflammatory signaling likely shapes the extent and directionality of microglia–neuron contacts observed during ICMS (Fig. 5F–H; Fig. 7F).

However, these observations remain correlative. To establish causality, future experiments should target specific pathways (e.g., P2Y12 knockout or pharmacological blockade) to test whether disrupting ATP/P2Y12 signaling decouples microglial process extension from neuronal activity during ICMS [141, 142]. Similarly, CX3CR1 knockout models could determine whether fractalkine-mediated chemotaxis underlies process polarization toward active neurons [143]. Such manipulations would reveal whether physical microglia–neuron contacts directly contribute to ICMS-induced neuronal adaptation or merely reflect underlying neuronal metabolic states. It is essential to investigate whether repeated, chronic 10 Hz ICMS maintains this homeostatic interplay or ultimately triggers maladaptive microglial states. If prolonged stimulation shifts microglia toward a pro-inflammatory phenotype, it could compromise long-term neuromodulation efficacy. Understanding these dynamics will be critical for designing stimulation protocols that sustain beneficial microglial engagement and avoid deleterious inflammation in chronic implant settings.

## Limitations

While 10-Hz ICMS directed microglial processes toward active neurons (Fig. 5F–H) and contact frequency correlated with neuronal adaptation (Fig. 7F; Fig. 8F), several caveats apply. First, we analyzed only soma contacts, omitting interactions with dendrites, axons, or synapses, where microglia prune weak connections and influence neurotransmission [96, 144]. Second, the one-hour ICMS window captures only acute responses (Fig. 3E–F). Chronic stimulation may induce delayed glial adaptations, such as phagocytosis of stressed neurons or process retraction, and shifts in neuronal profiles (Fig. 6D–E), which require longitudinal imaging. Third, examining only 10 Hz limits the understanding of how microglia respond to other frequencies and pulse parameters. Higher frequencies (40–100 Hz) can induce pro-inflammatory microglial changes [139, 140], while kilohertz-range waveforms may suppress reactivity. A systematic exploration of frequency, charge density, and waveform shape is needed to identify regimes that avoid unintended activation. Finally, although ATP/P2Y12 and fractalkine/CX3CR1 signaling likely mediate process extension toward active neurons [125, 126, 134, 138], we did not directly test these pathways. Blocking P2Y12 or CX3CR1 could reveal whether microglial motility requires those signals. We also did not measure microglial Ca²⁺ transients, which drive process movement[132], so it remains unclear if ICMS directly triggers intracellular signaling. Future studies should address these gaps to clarify how ICMS shapes microglial contributions to circuit modulation.

## Conclusion

This study demonstrates that low-frequency (10-Hz) ICMS does not provoke classical microglial activation but appears to engage microglia in a manner that correlates with neuronal activity. Through real-time imaging, we revealed that microglial processes selectively target active neurons during stimulation, with contact frequency inversely related to neuronal adaptation magnitude. These results suggest that microglia may play a homeostatic role in dampening overactive neurons and maintaining circuit stability. Crucially, these interactions were not driven solely by the electric field, but likely by neuron-derived signals, indicating that microglia are involved in shaping ICMS outcomes. Thus, microglia may contribute to the adaptations observed in neurostimulation techniques, including sensory neuroprostheses.

These findings offer important implications for neurostimulation technologies. By recognizing the potential for microglia to influence neuronal activity, future research could refine stimulation paradigms to optimize neuron-glia communication, potentially enhancing long-term stimulation efficacy while minimizing inflammatory responses. Longitudinal studies with targeted perturbations of microglia-neuron interactions would help clarify the causal role of microglia in these processes. Ultimately, incorporating a glia-centric perspective may pave the way for more adaptive and precise neurostimulation therapies, moving beyond acute efficacy to long-term circuit stability.

## Acknowledgements

Authors would like to thank Vanshika Singh for valuable feedback on the manuscript.

## Declarations

### Ethics approval and consent to participate

All animal care and procedures were performed with the approval of the University of Pittsburgh Institutional Animal Care and Use Committee and in accordance with regulations specified by the Division of Laboratory Animal Resources.

### Consent for publication

Not applicable

### Availability of data and materials

The datasets generated and used during the current study are available from the corresponding author on reasonable request.

### Author’s Contributions

KC, KS, and TK designed and performed experiments. CP and TK analyzed data and CP, KS, and TK wrote the manuscript.

*Acknowledgements*

## Data Availability

The raw/processed data required to reproduce these findings cannot be shared at this time as the data also forms part of an ongoing study

## Supplemental Materials

**Supplemental Figure 1.**
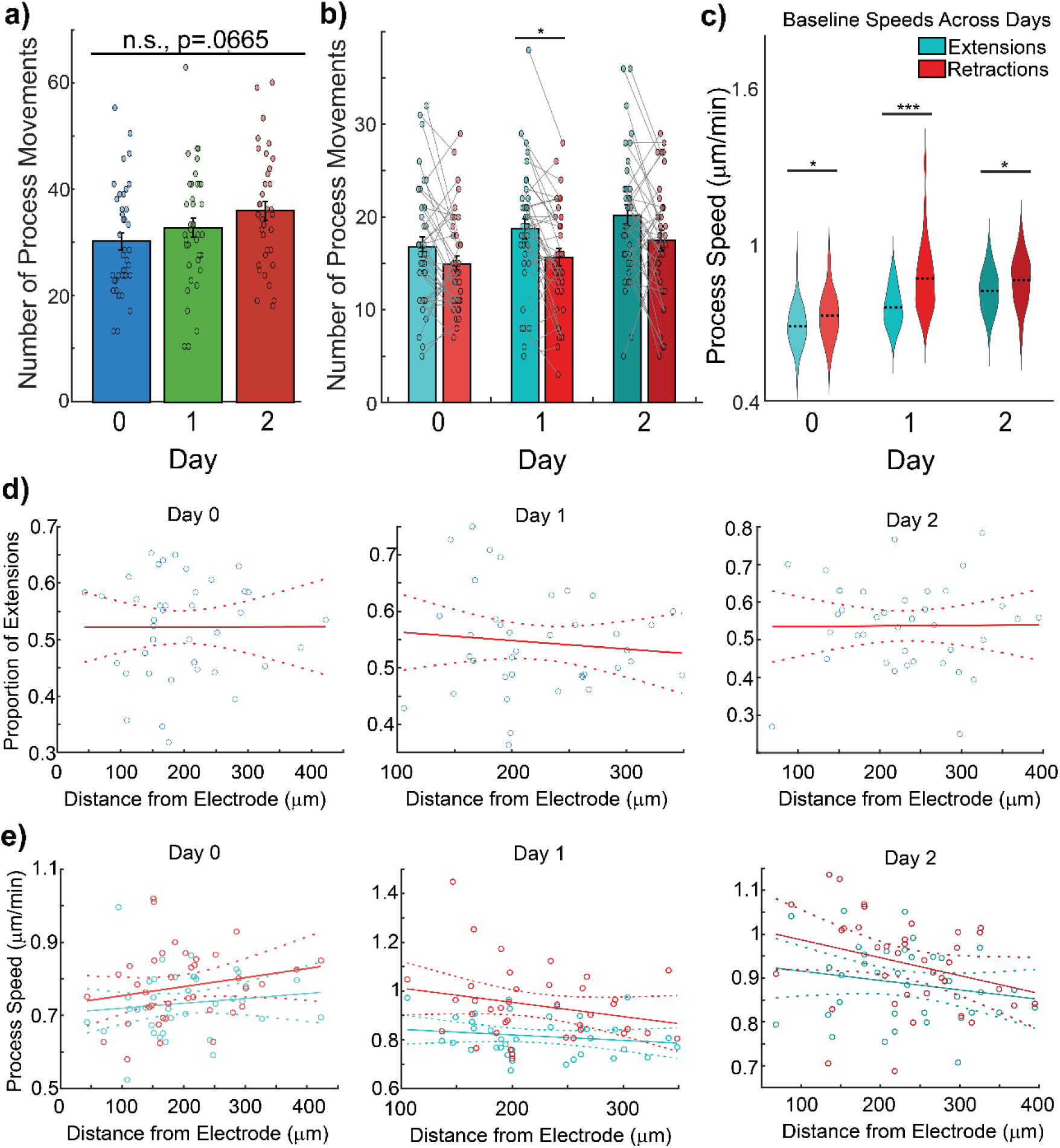
The balance of microglia process movements remains post-implantation, with retractions significantly faster than extensions. a) The total number of microglia process movements does not significantly increase with chronic microelectrode insertion (One-Way ANOVA, p = 0.067). b) There is a consistent trend of more extensions than retractions with only a significantly higher number of extensions 1-day post-implantation (One-Way ANOVA, p = .180, .038, .088). c) Process retractions are significantly faster than extensions for all days (One-Way ANOVA, p = .036, Kruskal-Wallis, p = 4.65e-05, One-Way ANOVA, p = .045). d) The proportion of extensions to retractions does not significantly change with microglia distance from the electrode regardless of the day (p = .991, .550, .957). e) Neither the speed of extensions nor retractions changes significantly as a function of distance from the electrode for any day post-implantation p = .442, .215, .312, .157, .250, .075).

**Supplemental Figure 2.**
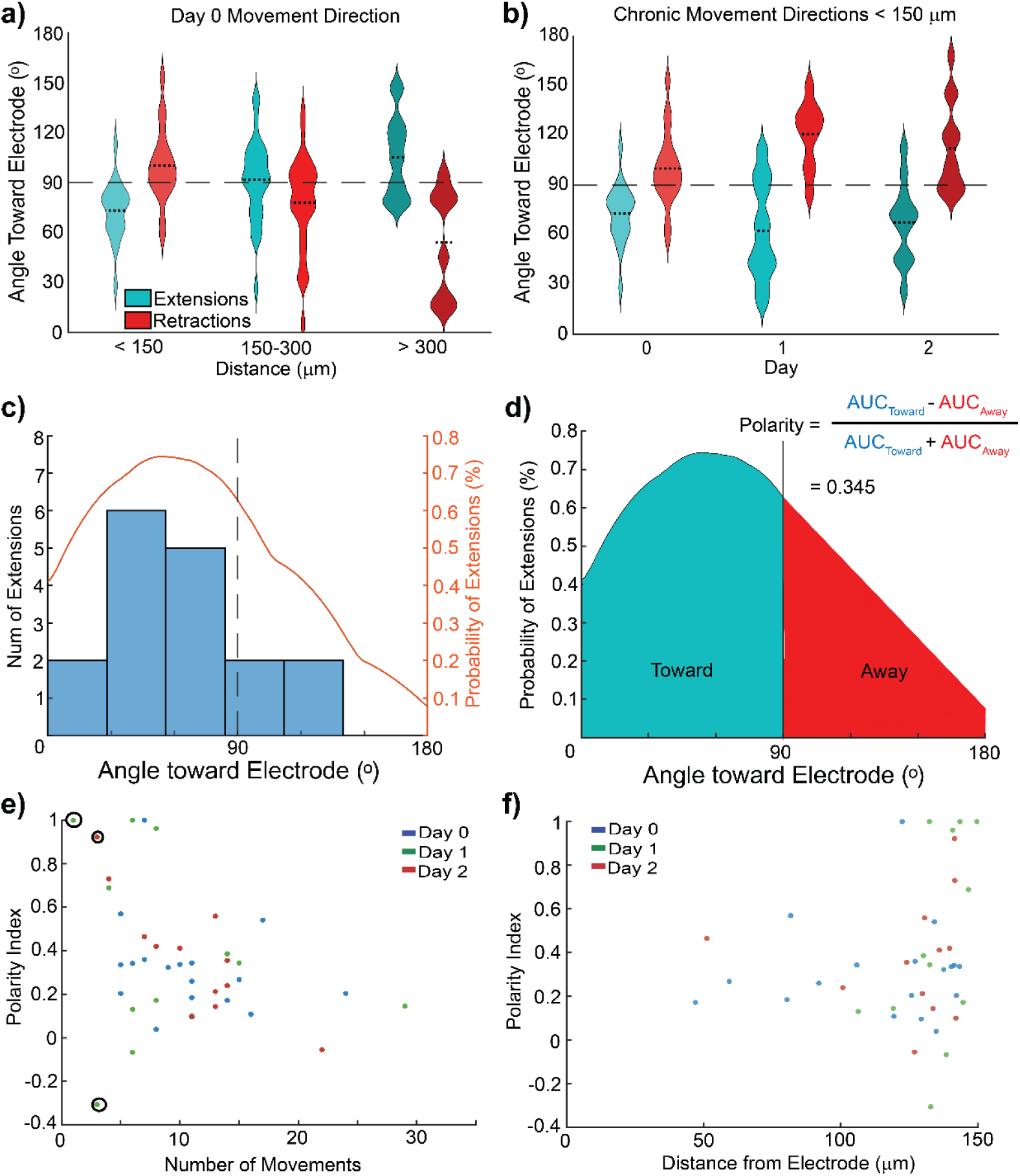
Microglial process retraction opposes extension: polarity index reflects dynamic balance during ICMS. a) The relative angles of extensions and retractions are significantly different and distributed in opposing directions, affirming that retractions typically occur along the same axes as extensions (One-Way ANOVA, p = .0002, .048, .0062). b) This opposition between extension and retraction directions holds true for microglia within 150 um of the electrode across days (One-Way ANOVA, p = .0002,3.72e-5, .0003). c) Example histogram of microglia extensions relative to the electrode and the resulting probability density function (PDF) calculated from the distribution. d) Example process for calculating the polarity index (p-index) using the PDF. e) Sample plot of polarity index compared to the number of movements, showing that variance is extreme when there are fewer than 4 movements and five potential values for the p-index. f) Plot of polarity index versus distance for microglia within 150 um of the electrode indicating increasing variance across days but no significant differences with distance.

**Supplemental Figure 3.**
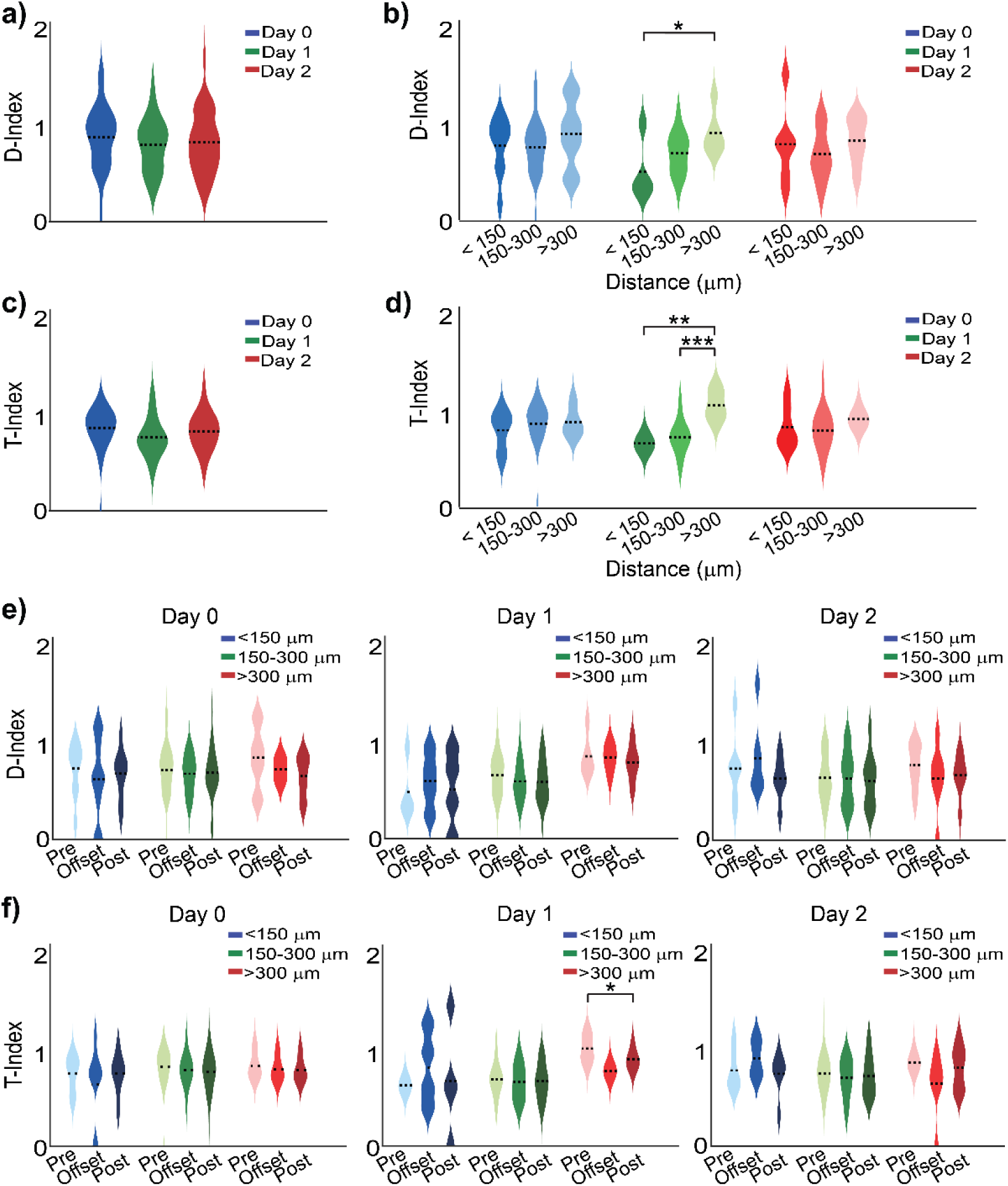
Microglia morphology changes across days according to distance from electrode implants regardless of stimulation. a) D-Index does not significantly change across days (One-Way ANOVA, p = .542). b) D-index is significantly lower on Day 1 within 150 µm of the electrode (One-Way ANOVA, p = .046). c) T-Index does not significantly change across days (One-Way ANOVA, p = .883). d) 10 Hz ICMS does not significantly modulate the length of microglia processes relative to the electrode and thus does not modulate another morphological indicator of microglia activation (One-Way ANOVA, p = .0034, .0007). e) D-Index does not significantly change with stimulation regardless of distance across days (One-Way ANOVA, p = .883, .351, .839). f) There was a significant decrease in T-Index in microglia more than 300 µm away from the electrode pre- and post-stimulation on Day 1 (One-Way ANOVA, p = 0.017).

**Supplemental Figure 4.**
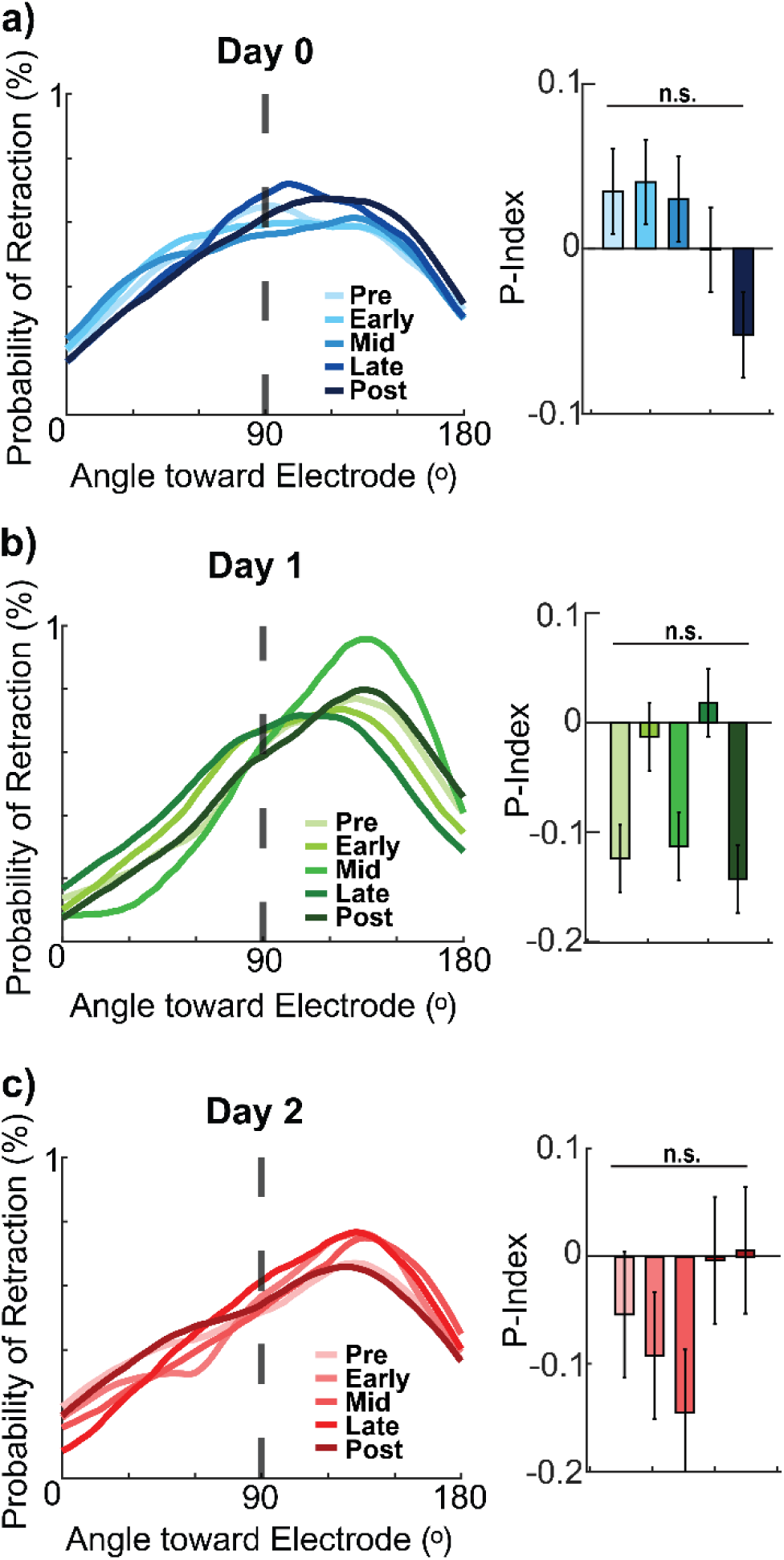
10 Hz ICMS does not significantly affect the direction of microglia retractions relative to the electrode. a) 10 Hz ICMS on Day 0 does not significantly modulate the direction of microglia process retractions nor the polarization of process retraction (Kolmogorov-Smirnov, p > .38 and Repeated Measures ANOVA, p = .713). b) Although Day 1 exhibits process retraction polarization away from the electrode, ICMS does not significantly modulate the directionality of movement, nor the polarity (Kolmogorov-Smirnov, p > 0.17 and Repeated Measures ANOVA, p = .228). c) Similarly, ICMS on Day 2 does not significantly drive microglia extensions toward the electrode nor affect polarization (Kolmogorov-Smirnov, p > 0.21 and Repeated Measures ANOVA, p = .293).

**Supplemental Figure 5.**
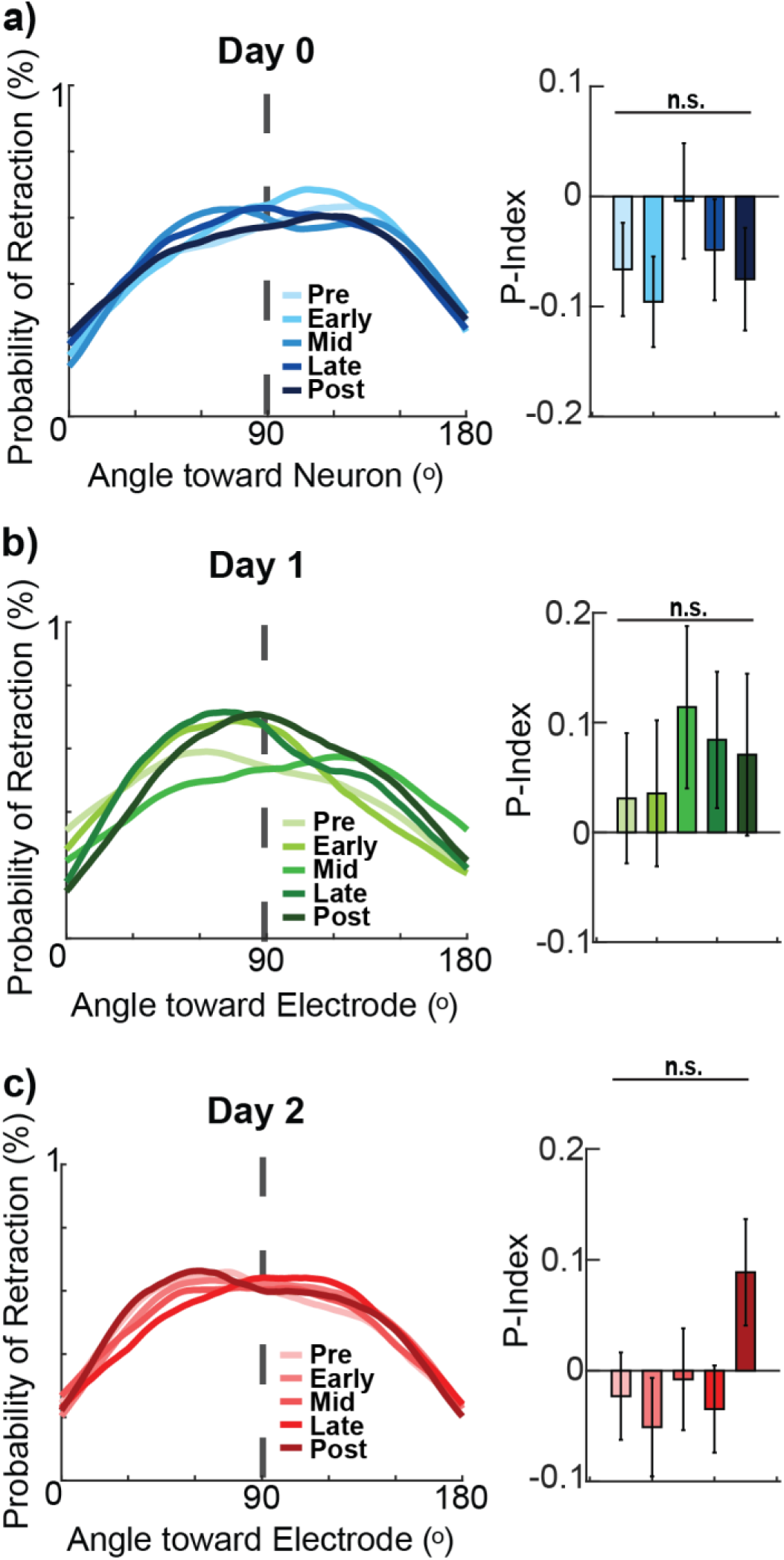
Microglia process extensions do not change near non-activated neurons. a) 10 Hz ICMS on Day 0 does not significantly modulate the direction of microglia process retractions nor the polarization of process retraction (Kolmogorov-Smirnov, p > .27 and Repeated Measures ANOVA, p = .406). b) Although Day 1 exhibits process retraction polarization away from the electrode, ICMS does not significantly modulate the directionality of movement, nor the polarity (Kolmogorov-Smirnov, p > 0.22 and Repeated Measures ANOVA, p = .149). c) Similarly, ICMS on Day 2 does not significantly drive microglia extensions toward the electrode nor affect polarization (Kolmogorov-Smirnov, p > 0.45 and Repeated Measures ANOVA, p = .310).

**Supplemental Figure 6.**
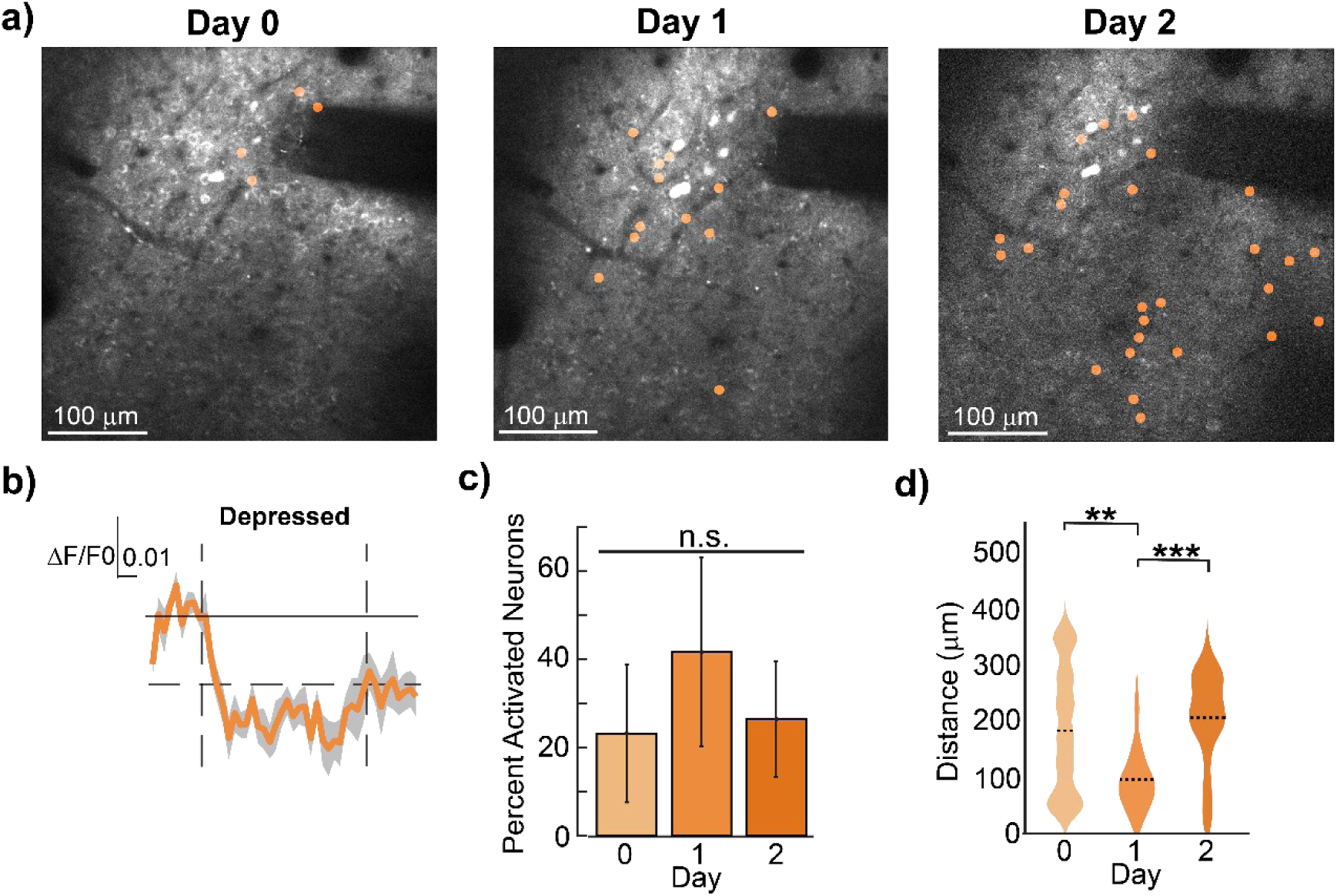
Spatial and temporal characterization of neurons exhibiting depressed activity during prolonged 10-Hz ICMS. a) Representative chronic images of neurons that exhibited depressed activity during 10 Hz ICMS. b) Representative calcium traces including stimulation onset and offset as well as the threshold for classification as depressed. c) The percentages of identified neuron profiles do not significantly vary across the days (Repeated Measures ANOVA, p = .788). d) Day 1 demonstrated depressed neurons significantly closer to the electrode site (Kruskal-Wallis, p = .0075, .510, 6.54e-10).

**Supplemental Figure 7.**
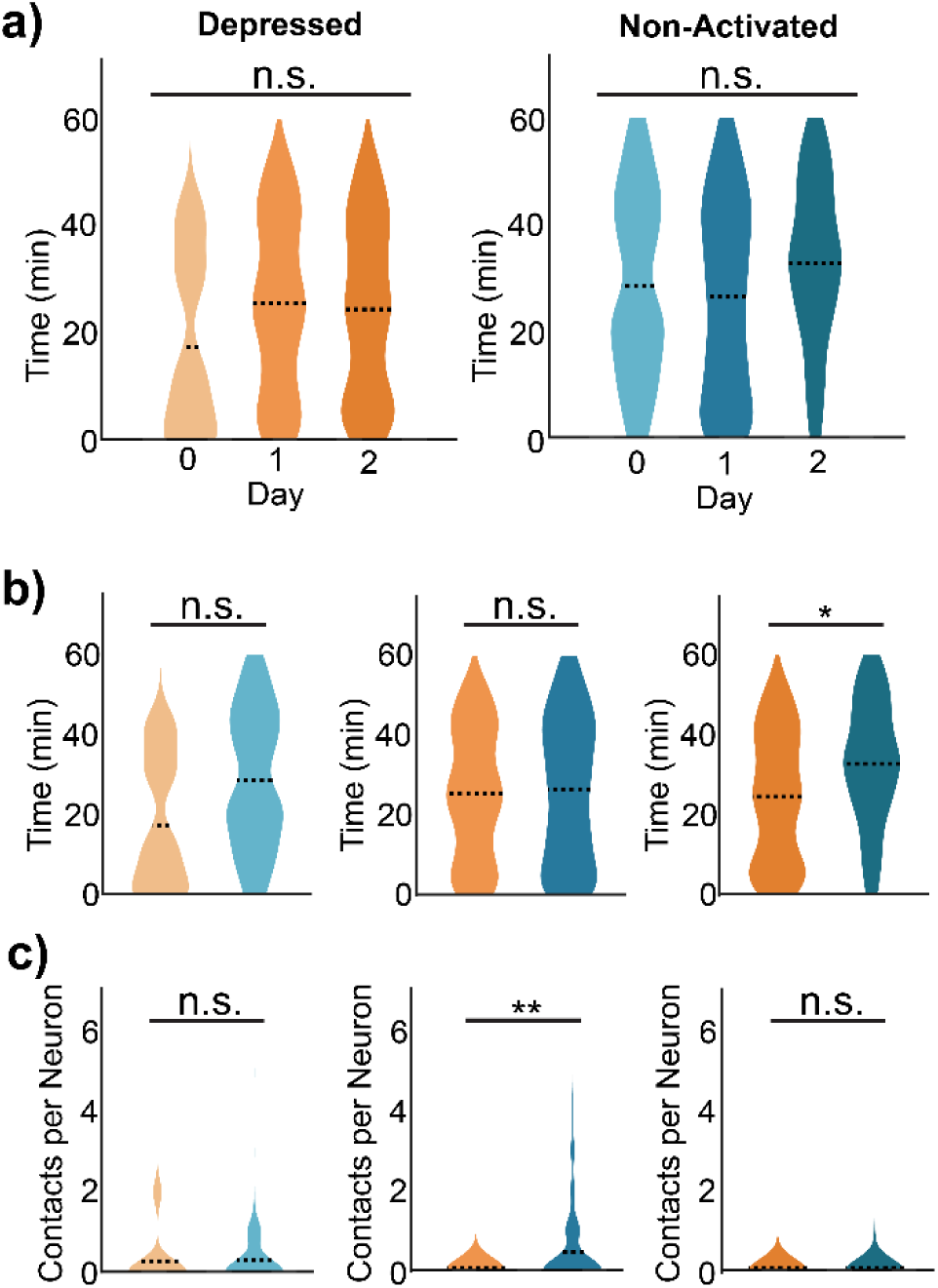
Microglia contact depressed neurons less frequently than non-activated neurons on Day 1. a)Peri-stimulation timing of microglia interactions with identified neuron activation profiles did not significantly differ across days (One-Way ANOVA, p = .617, Kruskal-Wallis, p = .101). b) Microglia contacted depressed neurons earlier into stimulation on Day 2, suggesting a potential dynamic homeostatic function during ICMS (One-Way ANOVA, p = .179, .826, .026). c) Microglia more frequently contacted non-activated neurons than depressed neurons on Day 2 (Kruskal-Wallis, p = .625, .006, One-Way ANOVA, p = .371).

