## Supplemental Materials for "Microglia surveillance is directed toward neuron activation during sustained intracortical microstimulation"

*** = denotes equal contributions**

**Affiliations:**

Supplemental Materials

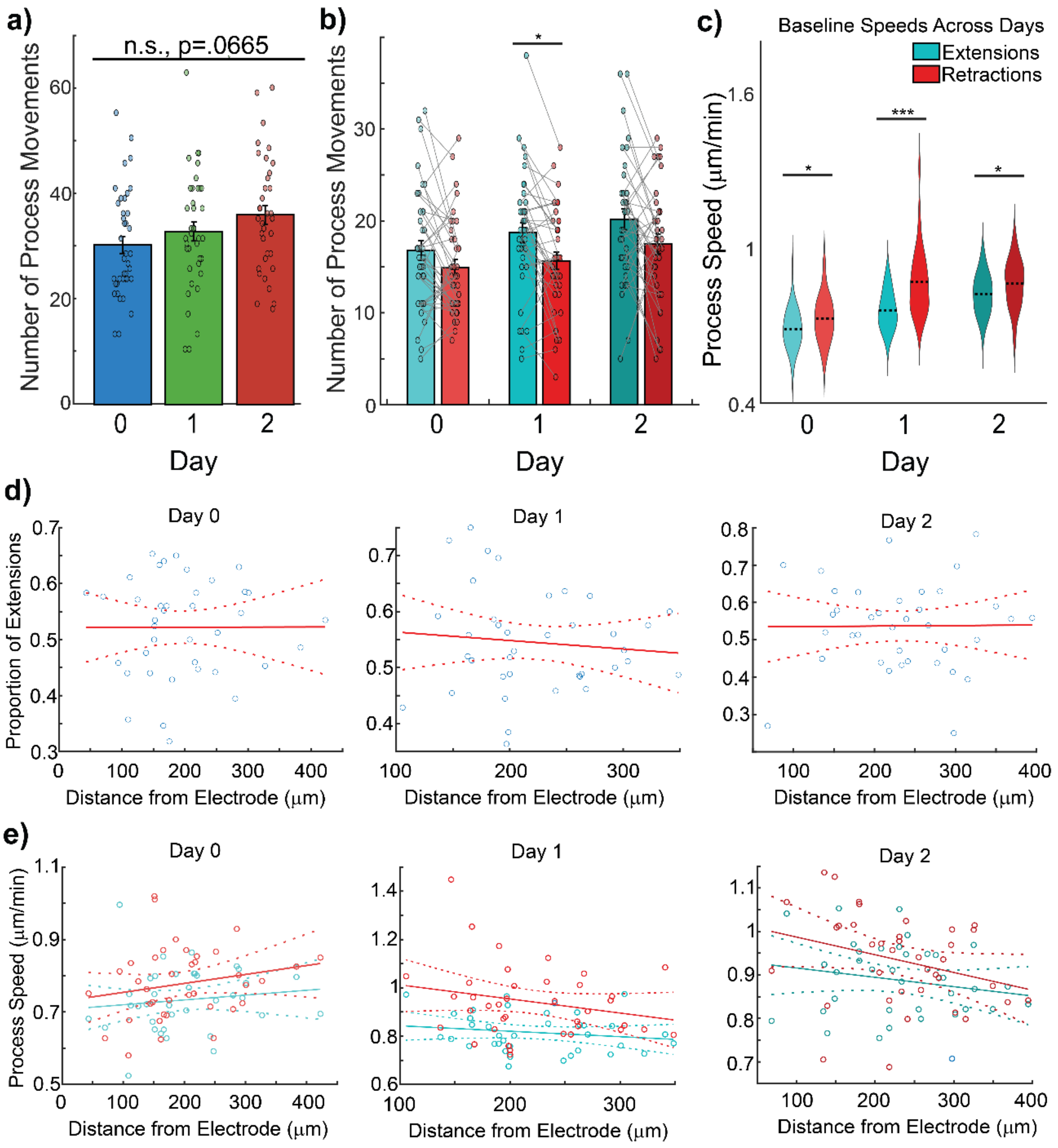


***Supplemental Figure 1. <u>The balance of microglia process movements remains post-implantation, with retractions significantly faster than extensions</u>.*** a) The total number of microglia process movements does not significantly increase with chronic microelectrode insertion (One-Way ANOVA, p = 0.067). b) There is a consistent trend of more extensions than retractions with only a significantly higher number of extensions 1-day post-implantation (One-Way ANOVA, p = .180, .038, .088). c) Process retractions are significantly faster than extensions for all days (One-Way ANOVA, p = .036, Kruskal-Wallis, p = 4.65e-05, One-Way ANOVA, p = .045). d) The proportion of extensions to retractions does not significantly change with microglia distance from the electrode regardless of the day (p = .991, .550, .957). e) Neither the speed of extensions nor retractions changes significantly as a function of distance from the electrode for any day post-implantation p = .442, .215, .312, .157, .250, .075).

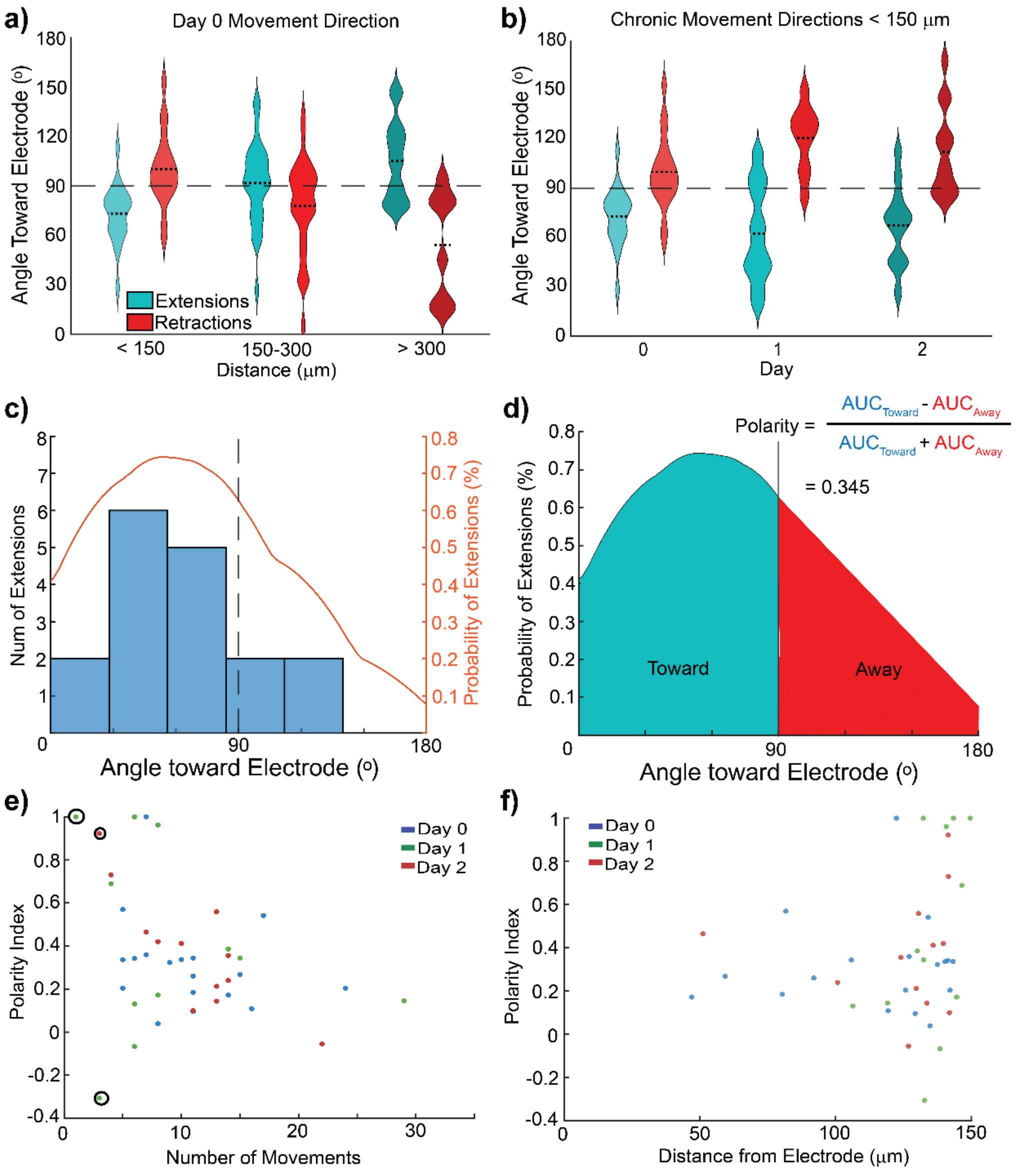


***Supplemental Figure 2. <u>Microglial process retraction opposes extension: polarity index reflects dynamic balance during ICMS</u>.*** a) The relative angles of extensions and retractions are significantly different and distributed in opposing directions, affirming that retractions typically occur along the same axes as extensions (One-Way ANOVA, p = .0002, .048, .0062). b) This opposition between extension and retraction directions holds true for microglia within 150 um of the electrode across days (One-Way ANOVA, p = .0002 ,3.72e-5, .0003). c) Example histogram of microglia extensions relative to the electrode and the resulting probability density function (PDF) calculated from the distribution. d) Example process for calculating the polarity index (p-index) using the PDF. e) Sample plot of polarity index compared to the number of movements, showing that variance is extreme when there are fewer than 4 movements and five potential values for the p-index. f) Plot of polarity index versus distance for microglia within 150 um of the electrode indicating increasing variance across days but no significant differences with distance.

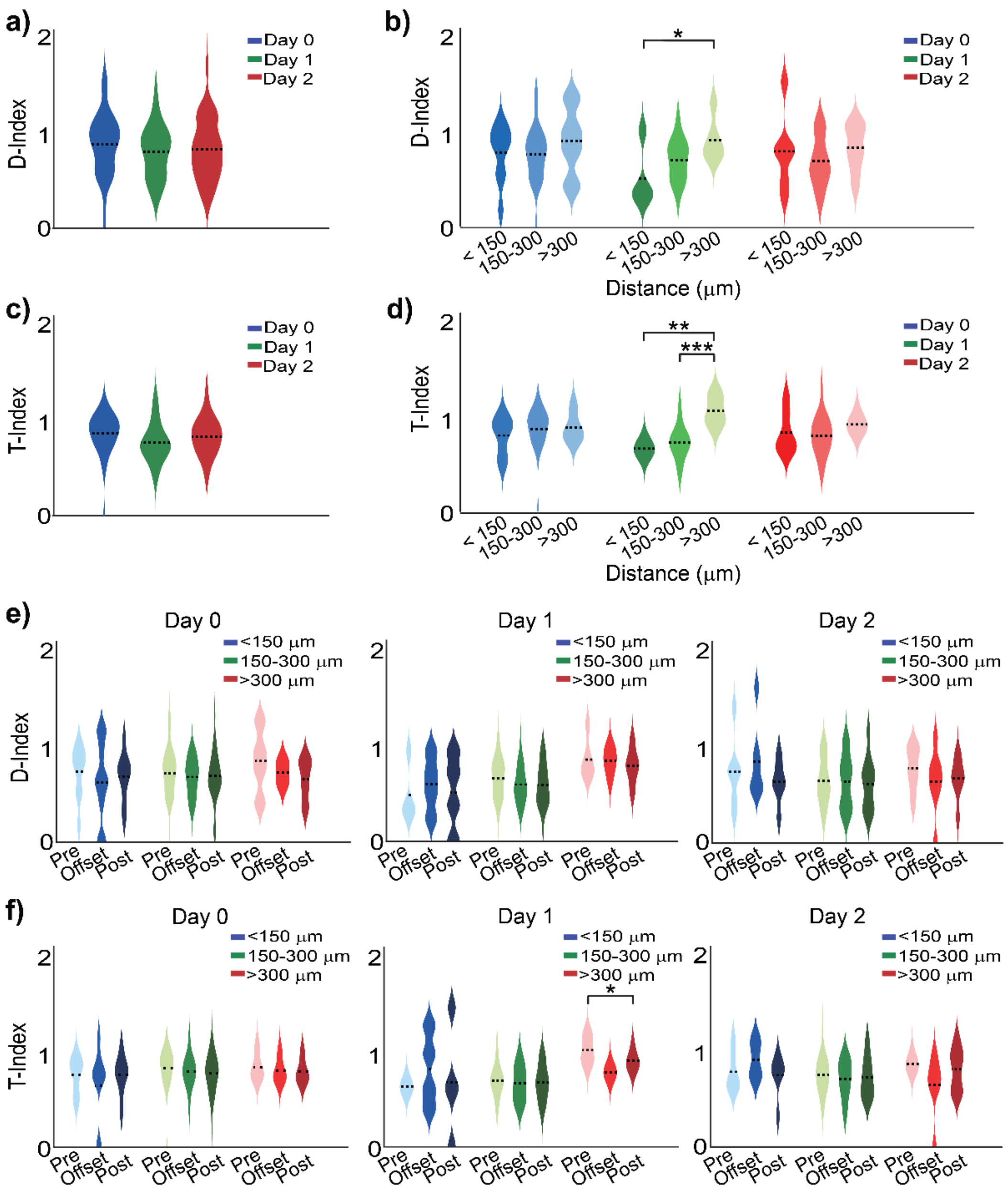


***Supplemental Figure 3. Microglia morphology changes across days according to distance from electrode implants regardless of stimulation.*** a) D-Index does not significantly change across days (One-Way ANOVA, p = .542). b) D-index is significantly lower on Day 1 within 150 μm of the electrode (One-Way ANOVA, p = .046). c) T-Index does not significantly change across days (One-Way ANOVA, p = .883). d) 10 Hz ICMS does not significantly modulate the length of microglia processes relative to the electrode and thus does not modulate another morphological indicator of microglia activation (One-Way ANOVA, p=.0034, .0007). e) D-Index does not significantly change with stimulation regardless of distance across days (One-Way ANOVA, p = .883, .351, .839). f) There was a significant decrease in T-Index in microglia more than 300 μm away from the electrode pre- and post-stimulation on Day 1 (One-Way ANOVA, p = 0.017).

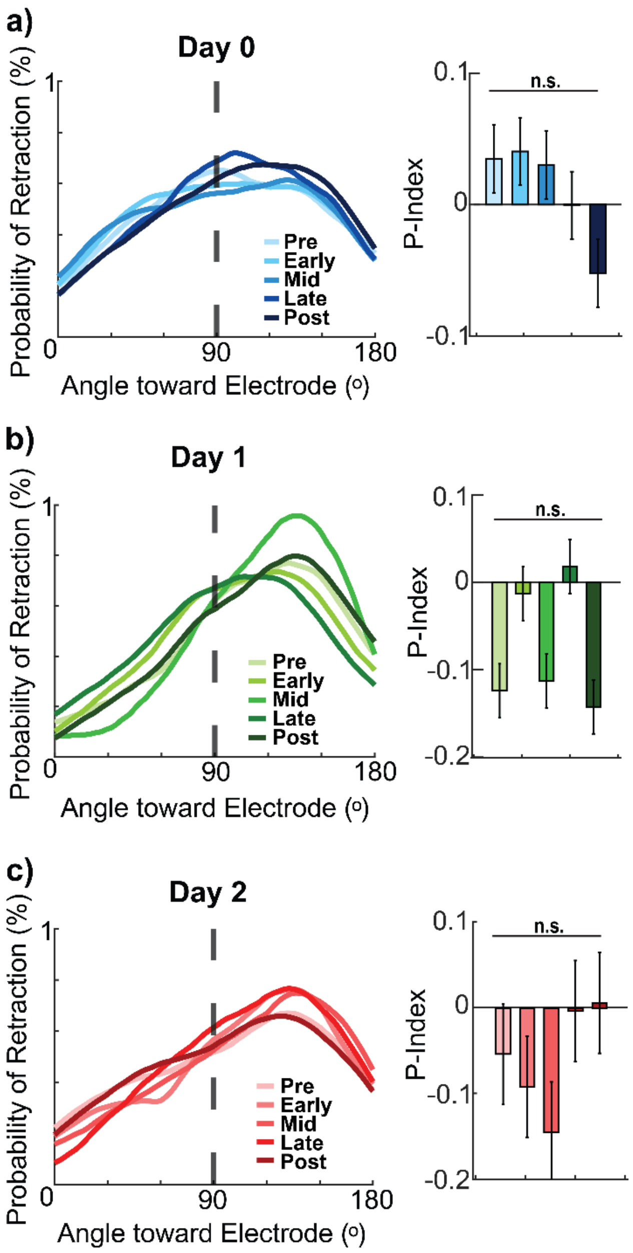


***Supplemental Figure 4.*** ***<u>10 Hz ICMS does not significantly affect the direction of microglia retractions relative to the electrode.</u>*** a) 10 Hz ICMS on Day 0 does not significantly modulate the direction of microglia process retractions nor the polarization of process retraction (Kolmogorov-Smirnov, $p > .38$ and Repeated Measures ANOVA, $p = .713$). b) Although Day 1 exhibits process retraction polarization away from the electrode, ICMS does not significantly modulate the directionality of movement, nor the polarity (Kolmogorov-Smirnov, $p > 0.17$ and Repeated Measures ANOVA, $p = .228$). c) Similarly, ICMS on Day 2 does not significantly drive microglia extensions toward the electrode nor affect polarization (Kolmogorov-Smirnov, $p > 0.21$ and Repeated Measures ANOVA, $p = .293$).

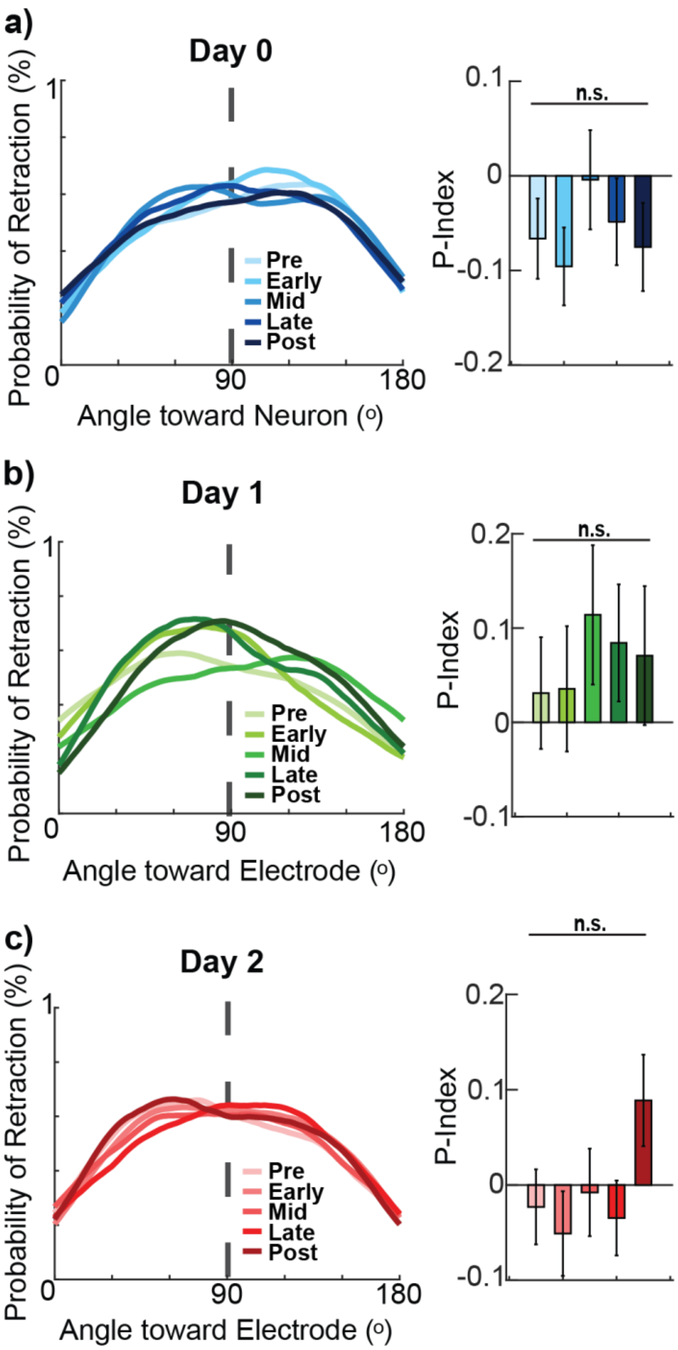


***Supplemental Figure 5. <u>Microglia process extensions do not change near non-activated neurons</u>.*** a) 10 Hz ICMS on Day 0 does not significantly modulate the direction of microglia process retractions nor the polarization of process retraction (Kolmogorov-Smirnov, $p > .27$ and Repeated Measures ANOVA, $p = .406$). b) Although Day 1 exhibits process retraction polarization away from the electrode, ICMS does not significantly modulate the directionality of movement, nor the polarity (Kolmogorov-Smirnov, $p > 0.22$ and Repeated Measures ANOVA, $p = .149$). c) Similarly, ICMS on Day 2 does not significantly drive microglia extensions toward the electrode nor affect polarization (Kolmogorov-Smirnov, $p > 0.45$ and Repeated Measures ANOVA, $p = .310$).

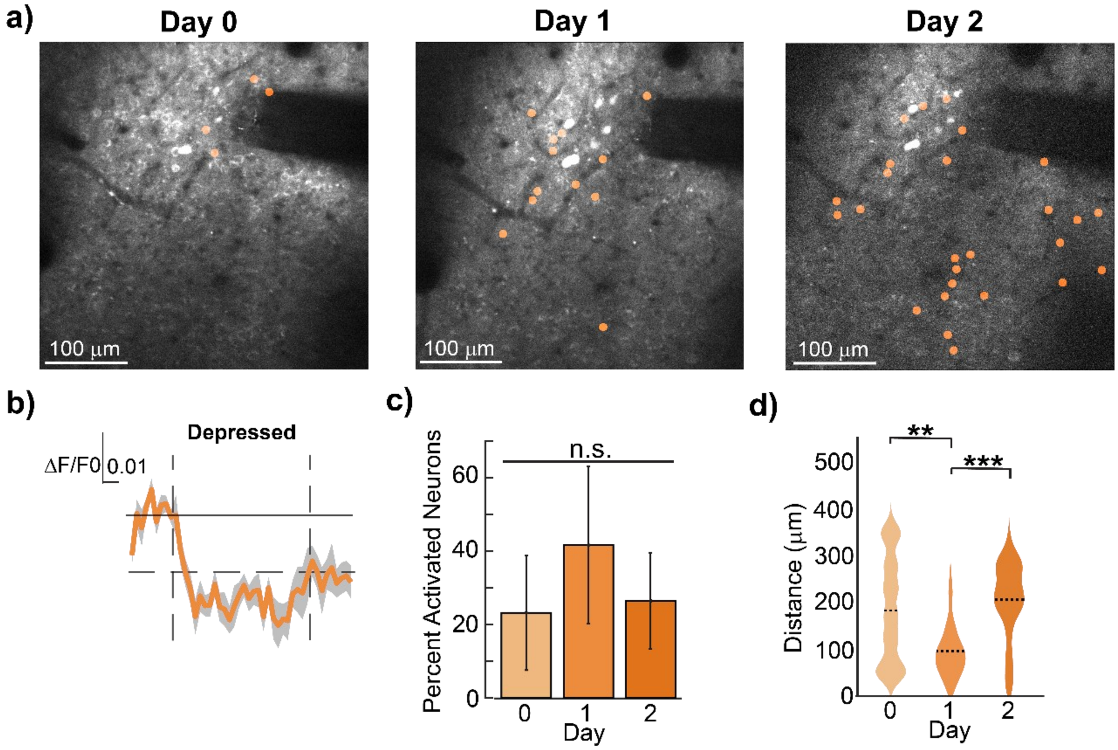


***Supplemental Figure 6. <u>Spatial and temporal characterization of neurons exhibiting depressed activity during prolonged 10-Hz ICMS.</u>*** a) Representative chronic images of neurons that exhibited depressed activity during 10 Hz ICMS. b) Representative calcium traces including stimulation onset and offset as well as the threshold for classification as depressed. c) The percentages of identified neuron profiles do not significantly vary across the days (Repeated Measures ANOVA, $p = .788$). d) Day 1 demonstrated depressed neurons significantly closer to the electrode site (Kruskal-Wallis, $p = .0075$, $.510$, $6.54e-10$).

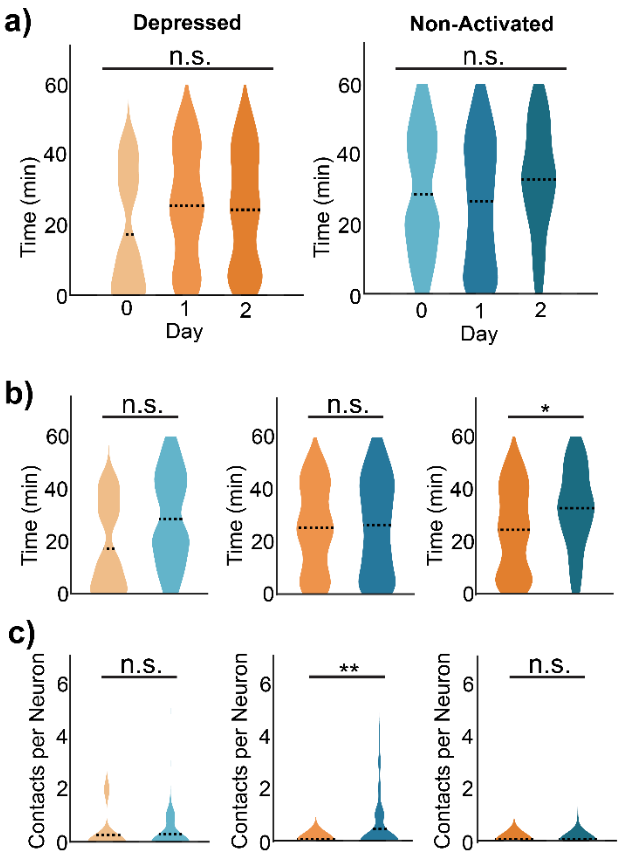


***Supplemental Figure 7. <u>Microglia contact depressed neurons less frequently than non-activated neurons on Day 1.</u>*** a) Peri-stimulation timing of microglia interactions with identified neuron activation profiles did not significantly differ across days (One-Way ANOVA, p = .617, Kruskal-Wallis, p = .101). b) Microglia contacted depressed neurons earlier into stimulation on Day 2, suggesting a potential dynamic homeostatic function during ICMS (One-Way ANOVA, p = .179, .826, .026). c) Microglia more frequently contacted non-activated neurons than depressed neurons on Day 2 (Kruskal-Wallis, p = .625, .006, One-Way ANOVA, p = .371).
